# Multimodal Analysis for Human ex vivo Studies Shows Extensive Molecular Changes from Delays in Blood Processing

**DOI:** 10.1101/2020.10.18.344663

**Authors:** Adam K. Savage, Miriam V. Gutschow, Tony Chiang, Kathy Henderson, Richard Green, Monica Chaudhari, Elliott Swanson, Alexander T. Heubeck, Nina Kondza, Kelli C. Burley, Palak C. Genge, Cara Lord, Tanja Smith, Zachary Thomson, Aldan Beaubien, Ed Johnson, Jeff Goldy, Hamid Bolouri, Jane H. Buckner, Paul Meijer, Ernest M. Coffey, Peter J. Skene, Troy R. Torgerson, Xiao-jun Li, Thomas F. Bumol

## Abstract

Multi-omic profiling of human peripheral blood is increasingly utilized to identify biomarkers and pathophysiologic mechanisms of disease. The importance of these platforms in clinical and translational studies led us to investigate the impact of delayed blood processing on the numbers and state of peripheral blood mononuclear cells (PBMC) and on the plasma proteome. Similar to previous studies, we show minimal effects of delayed processing on the numbers and general phenotype of PBMCs up to 18 hours. In contrast, profound changes in the single-cell transcriptome and composition of the plasma proteome become evident as early as 6 hours after blood draw. These reflect patterns of cellular activation across diverse cell types that lead to progressive distancing of the gene expression state and plasma proteome from native *in vivo* biology. Differences accumulating during an overnight rest (18 hours) could confound relevant biologic variance related to many underlying disease states.

## Introduction

Advances in “-omics” technologies now provide scientists with the ability to probe human biology and biologic variance with high sensitivity at the single cell level. These approaches have been of particular benefit to studies of immune-mediated diseases in humans where deep profiling of peripheral blood and peripheral immune cells has provided insights to underlying pathobiology, unique biomarkers of disease, and biological variation. The ability to reliably discern the state of individual cells and discover true biologic heterogeneity in a complex system like peripheral blood requires that the effects of sample handling and storage not overwhelm those associated with the underlying biology. The logistical challenges related to collecting, processing, and shipping blood in human studies and particularly in clinical trials can make the application of these highly sensitive technologies a challenge. At present, there is a dearth of multi-modal studies that provide practical guidance about how quickly peripheral blood samples need to be processed and which cells and cell pathways are most impacted by delayed processing.

Peripheral blood mononuclear cells (PBMC) are a workhorse of human immunology owing to the ease of collection and simplicity of cell isolation. Whole blood, from which PBMC are derived, is understood to be remarkably stable and previous work has established support for flexibility in sample handling for various whole blood assays, often as a way of managing the inherent challenges of human blood collection. Studies have shown that, when collected into anti-coagulant tubes to prevent clotting, whole blood left to “rest” was shown to be stable up to 24 hours at room temperature (Wu et al., 2017; Zini, 2014) and plasma profiling for common metabolites was similarly robust (van Eijsden et al., 2005; Zimmerman et al., 2012). Other data, however, demonstrated the potential for profound changes resulting from variability in sample handling, particularly in the context of untargeted assays such as those for transcriptomic analysis. Overnight delay of PBMC isolation from whole blood was shown to alter thousands of genes, in particular JUN, FOS, and the heat shock pathway (Baechler et al., 2004), and even delays as short as four hours led to substantial changes, especially in immune-related gene expression (Barnes et al., 2010; Massoni-Badosa et al., 2020). Particularly problematic are processing artifacts that impact a wide variety of genes and proteins related to immunity in health and disease, which can obscure the disease processes of interest (Dvinge et al., 2014). Missing from previous analyses, however, is a comprehensive, multi-modal approach from which to understand the full complexity of *ex vivo* biology and its impact on physiological signals.

In an effort to clearly address these questions prior to initiating a series of multi-center clinical studies focused on human immunology, we performed deep, multi-modal profiling of human peripheral blood stored in anticoagulant for varying lengths of time before processing to plasma and peripheral blood mononuclear cells (PBMC). This resource provides clear insights into the rapid changes related to delayed sample processing, elucidating the cells most altered, the cellular pathways most impacted, and the assays most affected. While flow cytometry did not reveal large-scale changes in cell type frequencies through an 18-hour delay, single-cell gene expression and high-plex plasma proteomics provide overwhelming evidence that cells of all types exhibit time-dependent changes that distorts the underlying biology. These changes are broad and dynamic, complicating the technical analysis of single-cell RNA-seq data and especially inferences of *in vivo* physiology from *ex vivo* assays. We propose the affected proteins and genes be carefully considered in any human biology study or clinical trial that uses blood and/or PBMCs to reflect *in vivo* biology, and that these findings may extend to blood- and immune-cell permeated tissue, as well. To assist in their use, we provide these data in an easily explorable web-accessible tool (http://bloodprocessingdelay.allenimmunology.org). The baseline cytometry, proteomic, and transcriptomic data on 10 donors serve as a high-quality resource to accelerate human systems immunology research and provide the substrate to begin decoding these effects in existing and emerging studies.

## Results

### Bulk transcriptomics identifies time-dependent changes unrecognized by cytometry

To study the effects of delays in PBMC processing from whole blood we performed two similar but independent experiments (Figure 1A). In Experiment 1, we isolated PBMC or plasma from whole blood at 2, 4, 6, 8, and 18 hours after blood draw from healthy donors (n=3) or those diagnosed with systemic lupus erythematosus (SLE, n=3). In Experiment 2, we assayed PBMC or plasma isolated from only healthy donors (n=4) starting at 2, 4, 6, 10, 14, and 18 hours after blood draw. In both experiments, the whole blood was held in the dark at room temperature prior to PBMC isolation by Ficoll gradient separation or plasma isolation. PBMC were assayed after freeze/thaw by flow cytometry and 10x Genomics single-cell RNA-sequencing, and the plasma by Olink proteomics. Details of the samples, the assays used in each experiment, and any deviations are available in the Methods. Important to data interpretation, the samples were held as whole blood in phlebotomy tubes prior to processing. Thus, the samples were a closed system, obviating the confounding effects of cellular migration and blood cell development as sources of time-dependent variability.

**Figure 1.**
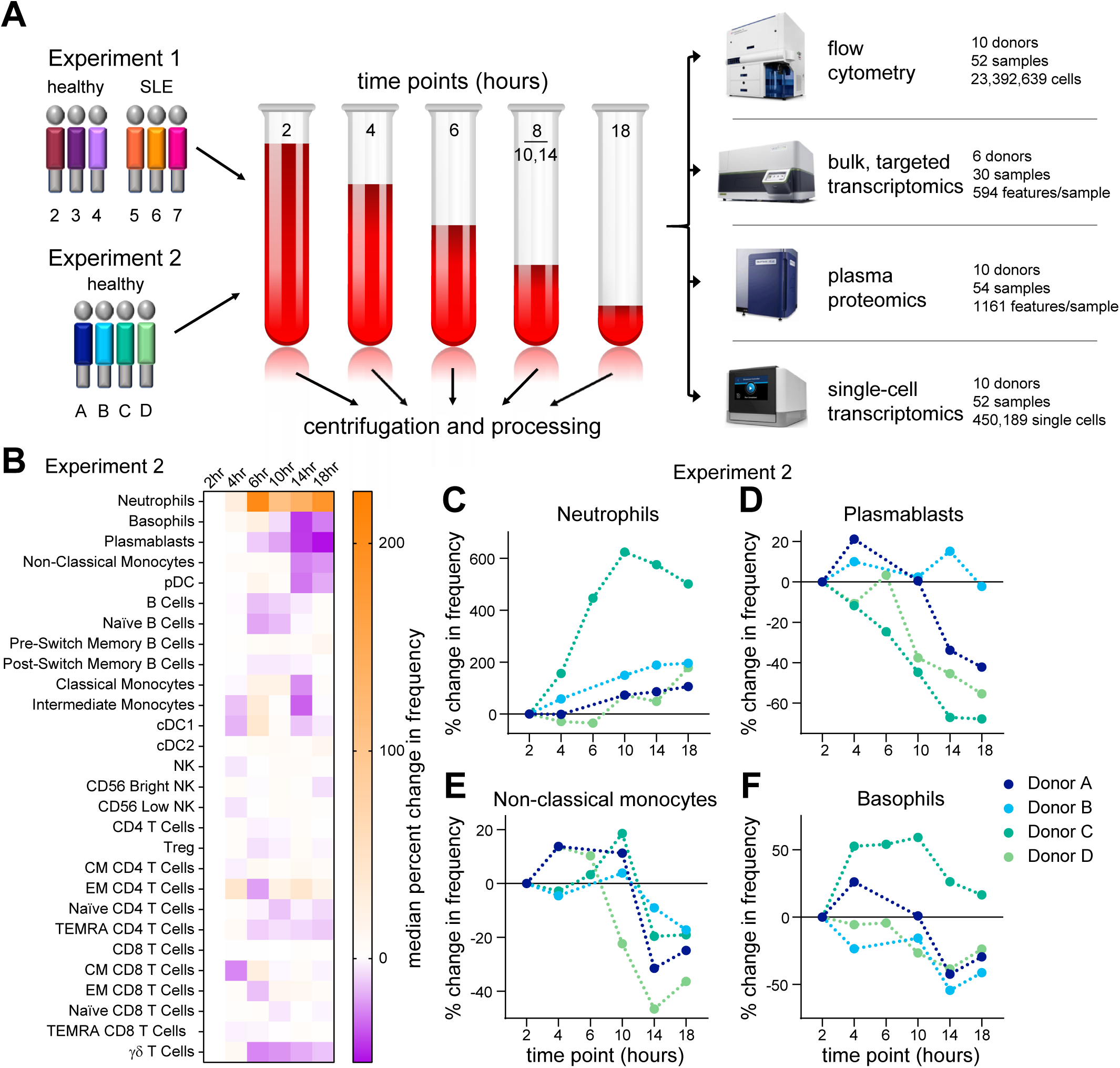
Flow cytometry suggests minimal effects from PBMC processing delay. (A) Schematic of the designs of Experiment 1 and Experiment 2. (B-F) Flow cytometry data from Experiment 2 were gated by traditional methods and the percent change in frequency was calculated for each population relative to the 2-hour PBMC processing time point. Because of technical artifacts, the 6-hour time points of donors A and B were excluded from analysis. (B) Heat map of the median percent change in frequency across the donors in Experiment 2. (C-F) The percent change in frequency for selected populations in individual donors as a function of time of PBMC processing post-blood draw. Data points connected by a line are from sample aliquots derived of the same blood draw pool. See also Supplemental Figures S1 and S2.

We started by assessing basic metrics from our technical processes. Cell yields from PBMC isolation, and their viability, did not exhibit time-dependent changes (**Supplemental Figures S1A-B**). Nor were the cells somehow more “fragile”, as they exhibited similar post-thaw recovery and viability at any time of isolation (**Supplemental Figures S1C-D**). This overall consistency was substantiated by flow cytometry profiling of the major immune cell types present in PBMC, most of which did not show noticeable changes throughout the processing delay time series (Figure 1B and **Supplemental Figure S2A-B**). Neutrophils (CD15^+^ SSC^hi^ cells), however, were apparently increased over the time course in both experiments (Figures 1C and **Supplemental Figure S2C-D**), consistent with previous data (McKenna et al., 2009; Nicholson et al., 1984), suggesting an increased recovery of low-density neutrophils by Ficoll gradient separation with longer delays in PBMC processing. In addition, more granular flow cytometry profiling in Experiment 2 showed decreases in plasmablasts, non-classical monocytes, and basophils (Figures 1D-F), though with the less extensive phenotyping in Experiment 1 we cannot confirm these observations. Nevertheless, with the exception of rare neutrophils, time-dependent effects on the abundance of cell types were modest.

The minor effects seen by cytometry could result from subtle changes in PBMC isolation and PBMC survival, but little can be inferred of cellular activation or stress from the flow cytometry panels used in this study to quantitate cell types. To get a better understanding of cellular activity, we first assayed bulk PBMC from the six donors in Experiment 1 by targeted transcriptomics with the Nanostring nCounter platform. This clinical assay enumerates a panel of 594 pre-selected immune- and disease-relevant transcripts from a bulk cell sample. Despite well documented person-to-person variability in PBMC gene expression (Kaczorowski et al., 2017; Radich et al., 2004; Whitney et al., 2003), Principal Component Analysis (PCA) indicated that PBMC processing delay accounted for a substantial part of the variability (**Supplemental Figure S3**). These effects were reflected in unsupervised clustering of the data, where it was apparent that expression levels across nearly the entire targeted gene set are inverted at 18 hours post-blood draw compared to the previous time point (Figure 2A, box indicated by the solid line). A feature of this longitudinal data set is that gene regulation dynamics can be observed, revealing that in all six donors a subset of these genes underwent an earlier induction before returning to low levels (box indicated by the dashed line). Aggregate analysis highlighted common and progressive exaggeration of changes over the time course (Figure 2B), with 85 genes significantly induced (including *IL20*, *DEFB4A*, *DUSP4*, *LILRB5*) and 261 genes significantly down-regulated (including PD-1 (*PDCD1*), ST2 (*IL1RL1*), *CX3CL1*, and *CCR2*) at 18 hours relative to 2 hours post-draw (Figure 2C-D and **Supplemental Table S1**). Without the context of the longitudinal data, these expression patterns could suggest physiologic biology that was not actually present in the host. Therefore, we conclude that while the cells themselves are generally robust to processing delays, their biological profile is greatly impacted.

**Figure 2.**
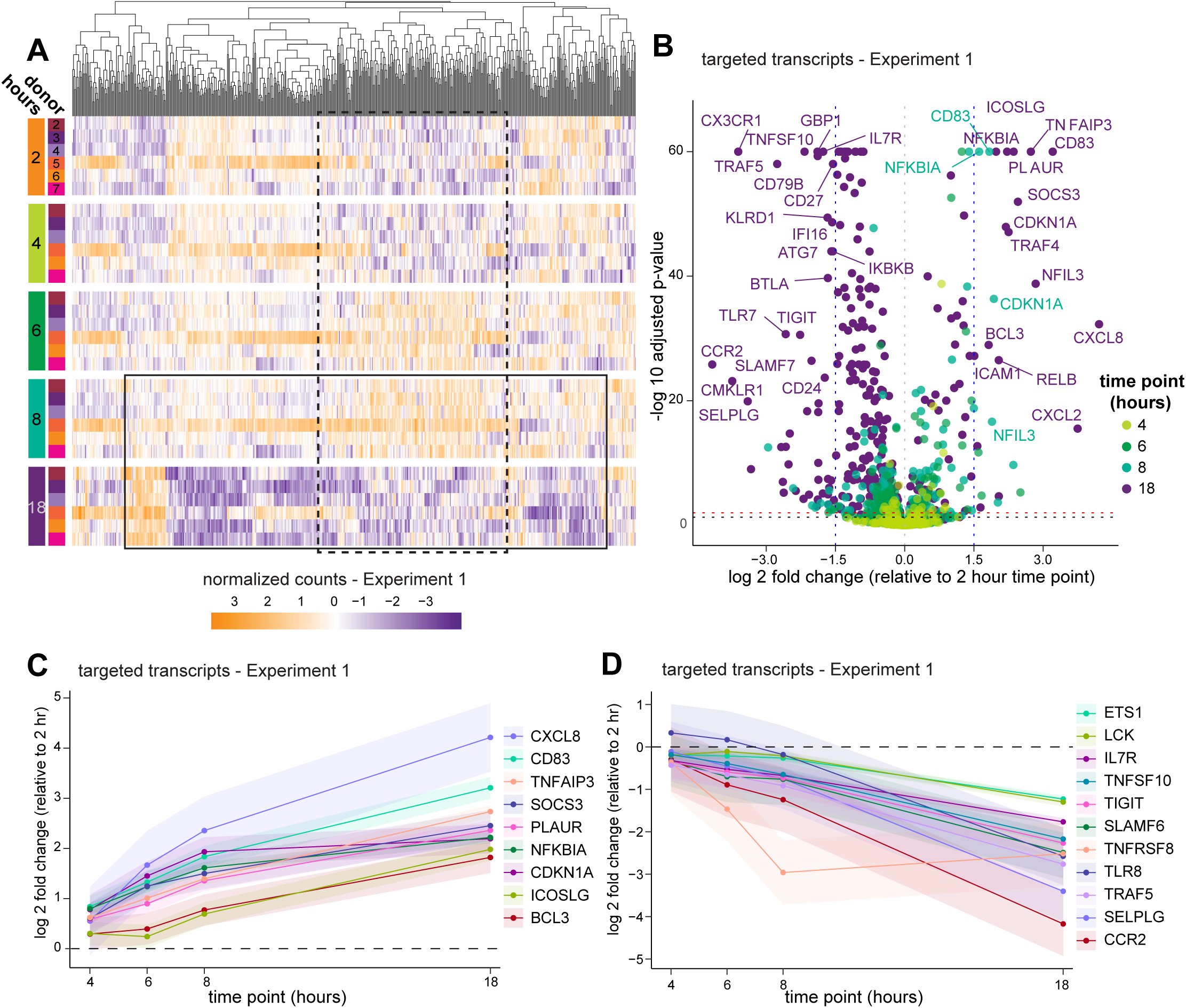
Bulk transcriptomics reveal profound changes in gene activity. PBMC in Experiment 1 were prepared from whole blood at various times after blood draw and assayed by targeted transcriptomics in bulk using Nanostring nCounter. (A) Targeted gene expression data colored by normalized counts relative to the mean of each gene across all samples. Genes (columns) are organized by unsupervised hierarchical clustering (dendrogram at top), and shown by time point (row groupings) post-blood draw and donor (rows). The boxes approximate subsets of gene that exhibit a pattern of inversion of expression between 8 and 18 hours post-blood draw (solid line) or an induction and re-normalization pattern (dashed line). (B) Volcano plot of all targeted gene expression transcripts in samples from Experiment 1 at each time point compared to 2 hours post-draw. Adjusted p-value cutoffs of 0.05 (red) and 0.01 (black), and log2 fold changes of |1.5| (blue) are indicated by dashed lines. A selection of the highest fold change transcripts are labeled with their gene names. (C-D) Data were analyzed using a generalized linear mixed effects model (details found in the Methods) and genes exhibiting significant change across the time course were selected for a similar and consistent pattern of up-regulation (C) or down-regulation (D). Shaded areas around the line indicate the 95% confidence interval. Changes are significant where the 95% CI does not include zero (dashed line). See also Supplemental Figure S3 and Supplemental Table S1.

### Delayed processing results in substantial changes to the plasma proteome

The extensive changes recognized by bulk transcriptomics suggested that whole blood awaiting processing was an active and dynamic immune environment, and for numerous reasons we hypothesized there would be significant changes in plasma proteins as blood awaited processing: it has previously been reported that storage of whole blood led to increased levels of thrombospondin by eight hours (Kaisar et al., 2016), immune cells are known to respond to hemostasis (Jenne et al., 2013), temperature (Cho et al., 2010) and oxygen levels (Nizet and Johnson, 2009), and many transcripts demonstrating increases in our bulk transcriptomics assay encode secreted proteins (**Supplemental Table S1**). We speculated that not only were cells likely undergoing an intrinsic response, but that the extrinsic environment was likely changing as well.

To test this hypothesis, we interrogated the 54 samples in our study for 1161 plasma proteins using a dual-antibody targeted assay (Assarsson et al., 2014). We found nearly one-third of the proteins showed relative abundance varying with delayed processing in a combined analysis of both experiments (Figure 3A and **Supplemental Table S2**). While the number of proteins exhibiting significant change was only modestly increased over the time course (350 proteins at 4 hours to 469 proteins at 18 hours post-draw), the magnitude of the changes exhibited a shift from 6 hours onward. Only 5 proteins were 1.5-fold changed between 2 and 4 hours post-draw (LAT, CD40L, EGF, PDGFβ, and SDC4), while by 6 hours 27 proteins had changed by more than 1.5-fold and by 18 hours 69 proteins had changed by at least 1.5-fold, relative to the 2-hour time point. Given that whole blood waiting in phlebotomy tubes is a closed system, increases in plasma proteins must be due to *de novo* production or enhanced availability to detection. The latter may result from liberation of sequestration factors or modification of assay-targeted epitopes. In all cases, increases in detectable proteins over time suggest active biology in the waiting blood.

**Figure 3.**
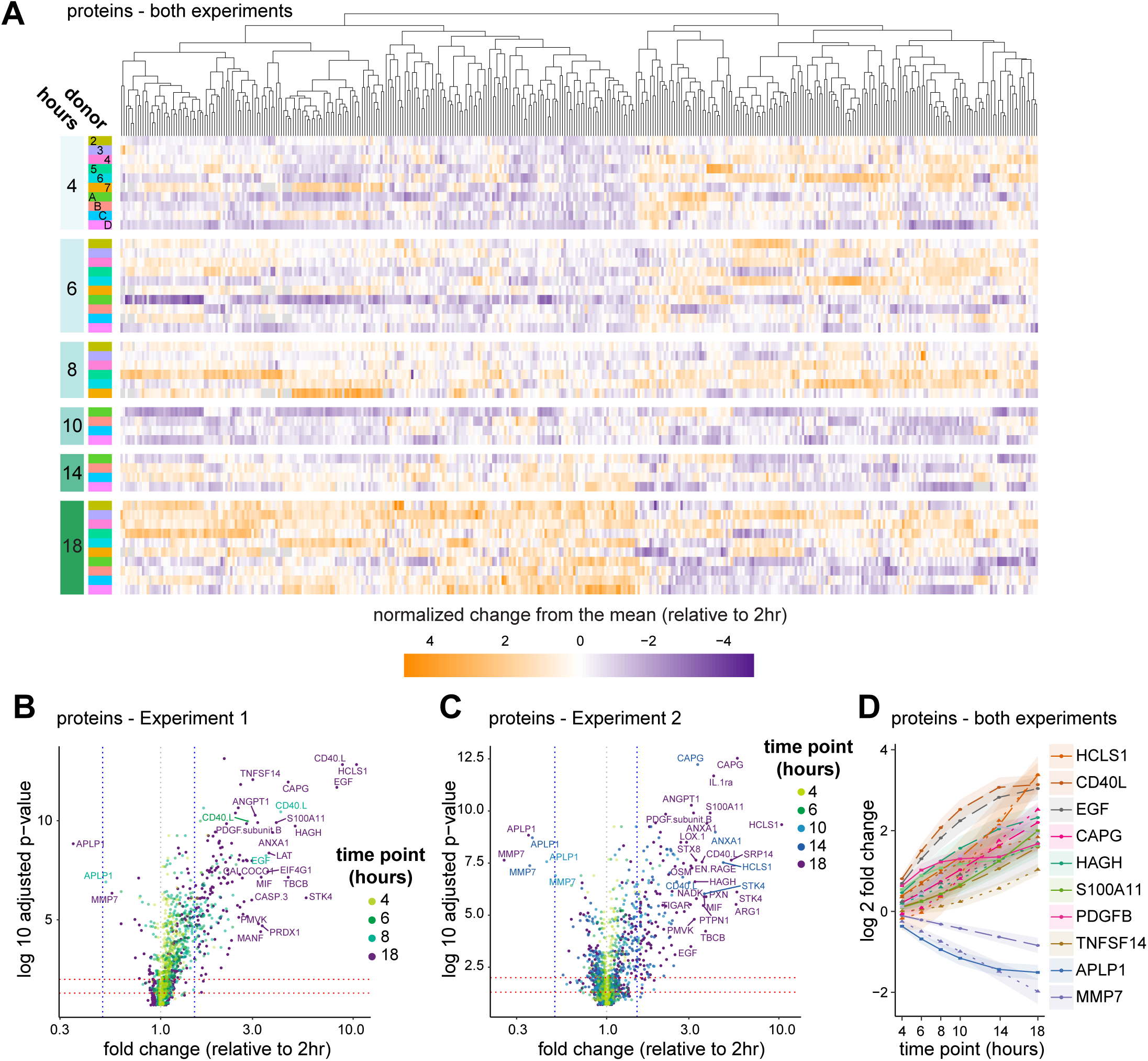
The plasma proteome reflects platelet and immune activation. Plasma proteins were quantitated by a dual-antibody proximity extension assay (Olink). (A) Data from all 10 donors were combined for analysis using a generalized linear mixed effects model (GLMEM) and only proteins exhibiting significant change are shown. Proteins (columns) are organized by unsupervised hierarchical clustering (dendrogram at top), shown by time point (row groupings) post-blood draw and donor (rows), and colored by normalized change from the mean for each time point relative to the 2-hour time point. (B-C) All proteins in Experiment 1 (B) and Experiment 2 (C) were plotted for fold change and adjusted p-value (Benjamini-Hochberg FDR) as calculated using a GLMEM model, and colored by time point. (D) A selection of proteins showing the greatest fold change in combined analysis were plotted by time point and shaded regions indicate the 95% confidence interval. Where both experiments identified a significant fold change for a given protein but those fold changes differed between experiments according to the GLMEM model, two lines are shown (dashed for Experiment 1, dotted for Experiment 2); otherwise a single line (solid) representing data from both experiments is shown. See also Supplemental Table S2.

Some of the most highly affected protein levels suggest complex and coordinated activity. For example, AnxA1 and S100A11 directly interact on early endosomes, and EGFR can both lead to phosphorylation of and be degraded by AnxA1 (Poeter et al., 2013). ANXA1, S100A11, and EGF protein are all increased in the plasma over time. More striking was that the number of platelet-related proteins significantly changed over time. In agreement with previous work (Khan et al., 2006), one of the earliest and most increased proteins was CD40LG (Figures 3B-D and **Supplemental Table S2**), a product of platelet activation Henn et al., 1998). PDGFβ, ANGPT1, STK4, STX8, OSM, and SDC4, were also rapidly and significantly increased. These proteins, some of which were reported previously (Shen et al., 2018), are all known to be involved in platelet biology (Andrae et al., 2008; Beck et al., 2017; Golebiewska et al., 2015; Londin et al., 2014; Tanaka et al., 2003), perhaps indicative of regulation or dysregulation of platelet activation in the waiting blood despite the presence of anti-coagulant. Fewer proteins exhibited decreases in abundance, with APLP1 the most striking. APLP1 can be processed by gamma-secretase but, unlike the homologs APP and APLP2, it was suggested that cleavage of APLP1 could occur without ectodomain shedding (Schauenburg et al., 2018), potentially sequestering the protein intracellularly. MMP7 also exhibited a significant decrease over the time course and was shown to be a target of platelet activation (Yang et al., 2020) and platelet-derived CXCL4 (Erbel et al., 2015). These data demonstrate that many of the strongest changes in plasma proteins in whole blood *ex vivo* relate to platelet biology.

### Single-cell transcriptomic profiling reveals reorganization of cell type-specific gene expression

While the majority of cell-type frequencies were essentially stable throughout the blood processing delay (Figure 1), plasma proteomics showed the signals available to those cells were dynamic and the corresponding bulk transcriptomics indicated similar changes in gene regulation, as might be expected from an immune system evolved to respond to extracellular signals. To better understand the impact of blood processing delay on distinct cell types we performed droplet-based single-cell RNA-sequencing on both experiments. In total, we collected and analyzed 450,189 high quality singlet cells with an average median features per cell of 1942 (**Supplemental Figure S4** and **Supplemental Table S3**), for a total of more than 800 million unique RNA molecules quantitated by single-cell RNA sequencing.

We assessed the overall effect of delayed blood processing across all cell types by aggregating together the samples in each experiment and summarizing the high-dimensional data set in two dimensions using tSNE (Figure 4 and **Supplemental Figure S5**). With this visualization, it was clear that by 18 hours post-draw (purple) there was a profound shift in the global gene expression pattern per cell across multiple clusters, with the time effect dominating inter-donor variability. While this effect was clear and pervasive at 18 hours in Experiment 1, we were concerned that the irregular time spacing (PBMC isolation at 2, 4, 6, 8, and 18 hours post-draw) was a confounder in the way the data clustered. We therefore changed the experimental design of Experiment 2 (PBMC isolation at 2, 4, 6, 10, 14, and 18 hours post-draw) so the later time points occurred at more regular intervals. The time-dependent divergence from the native transcriptional state was even more pronounced in Experiment 2, with a progressive distancing apparent at 10 hours (light blue), 14 hours (dark blue), and 18 hours (purple) post-draw. These effects were not restricted to a limited number of cell types, such as monocytes and dendritic cells (DC), but were apparent in all labeled cell types, even those thought to be quiescent, such as naive T cells (Sprent and Surh, 2011). These data demonstrate that analysis of global gene expression profiles in human PBMC can be confounded by pre-analytical process artifacts.

**Figure 4.**
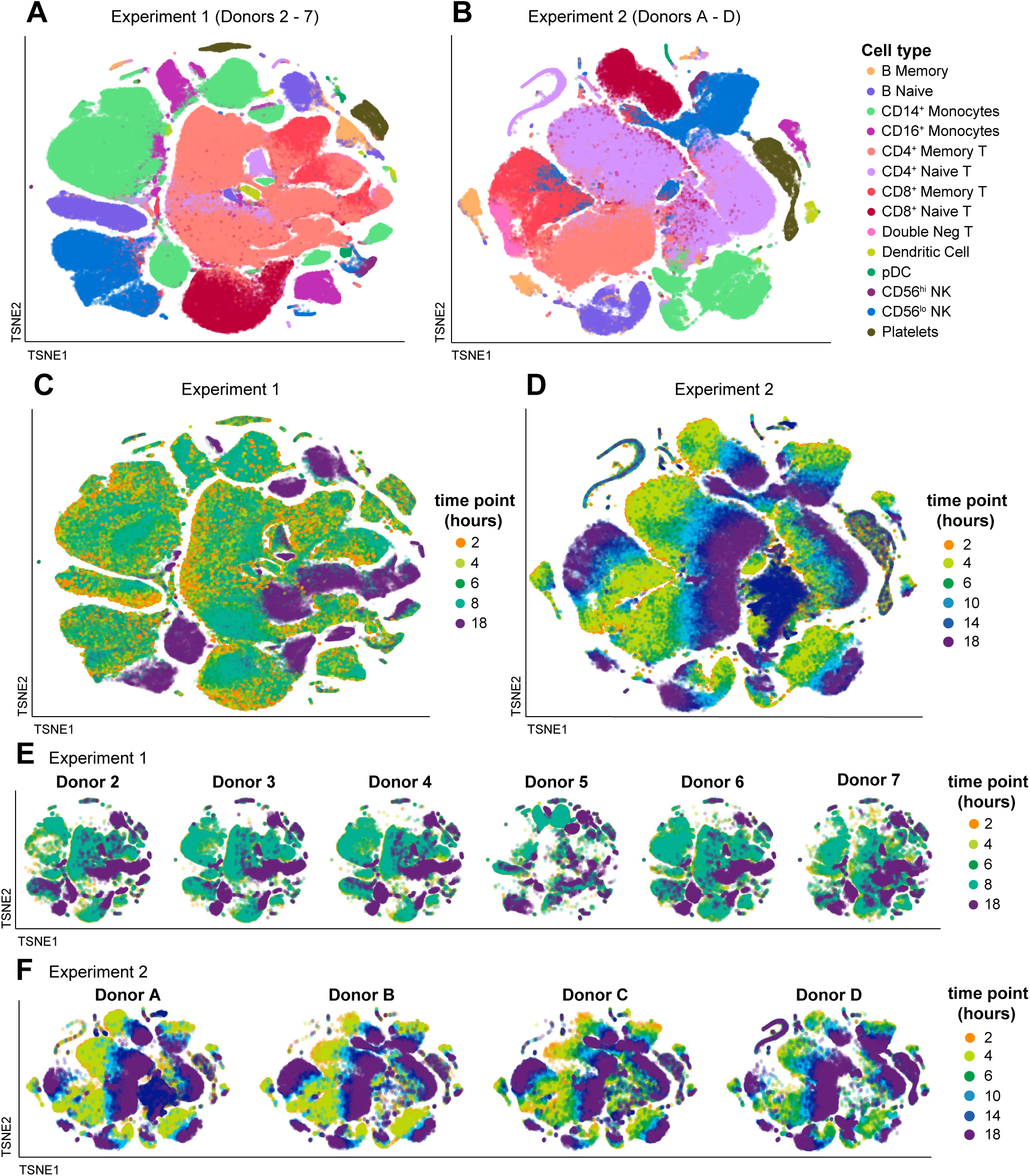
PBMC exhibit severe time-dependent changes after blood draw. The single-cell RNA-sequencing normalized gene expression matrices from each sample were identified for donor and time point and aggregated per experiment (Experiment 1: donors 2-7; Experiment 2: donors A-D). The multidimensional data was displayed in two dimensions using tSNE, colored as indicated. The data are overlaid in sequence with the latest time point (18hr) on top, obscuring some data points. Additional details can be found in the Methods. (A-B) tSNE visualizations colored by cell type for Experiment 1 (A) and Experiment 2 (B). Individual cells were assigned a cell type label independently of clustering or tSNE visualization. (C-F) tSNE visualizations colored by time point for Experiment 1 (C) and Experiment 2 (D). The tSNE maps in (C) and (D) were split out by donor for Experiment 1 (E) and Experiment 2 (F). See also Supplemental Figures S4 and S5, and Supplemental Table S3.

We were interested to more deeply understand the effect of blood processing delay on individual cell types to better define the changes on functionally distinct immune cell populations. To highlight cell type specific changes, we isolated the tSNE coordinates of the clusters mapping to selected cell types (Figures 5A) and quantified the occupancy of each cluster over the time course. In this approach, we infer that each cluster represents a distinct transcriptional state of a given cell type, such that a change in cluster composition within a cell type indicates changing transcriptional states. For example, most CD14^+^ monocytes start from a single, dense cluster at the earliest time point which disintegrates concurrently with the appearance of three new clusters by 18 hours, with the major transition occurring between 4 and 6 hours post-blood draw (Figure 5B). A similar pattern was observed for CD4^+^ memory T cells, CD8^+^ memory T cells, and double-negative T cells. In contrast, the originating clusters of CD8^+^ naive T cells and CD56^low^ NK cells persisted throughout the time course despite the appearance of new clusters, and CD4^+^ naive T cells exhibited some clusters persisting, some disappearing, and some appearing throughout the time course. Though the effects on different cell types resulted in different patterns of changing transcriptional profiles, for nearly all cell types changes were most apparent between the 4-hour and 6-hour time points (Figure 5C). This was true of cell types thought to be most responsive to environmental signals (e.g. monocytes) and for those thought to be quiescent (e.g. CD4^+^ naive T). From these data we conclude that measurements obtained from PBMC processed more than four hours after blood draw are increasingly at risk for pre-analytical artifacts.

**Figure 5.**
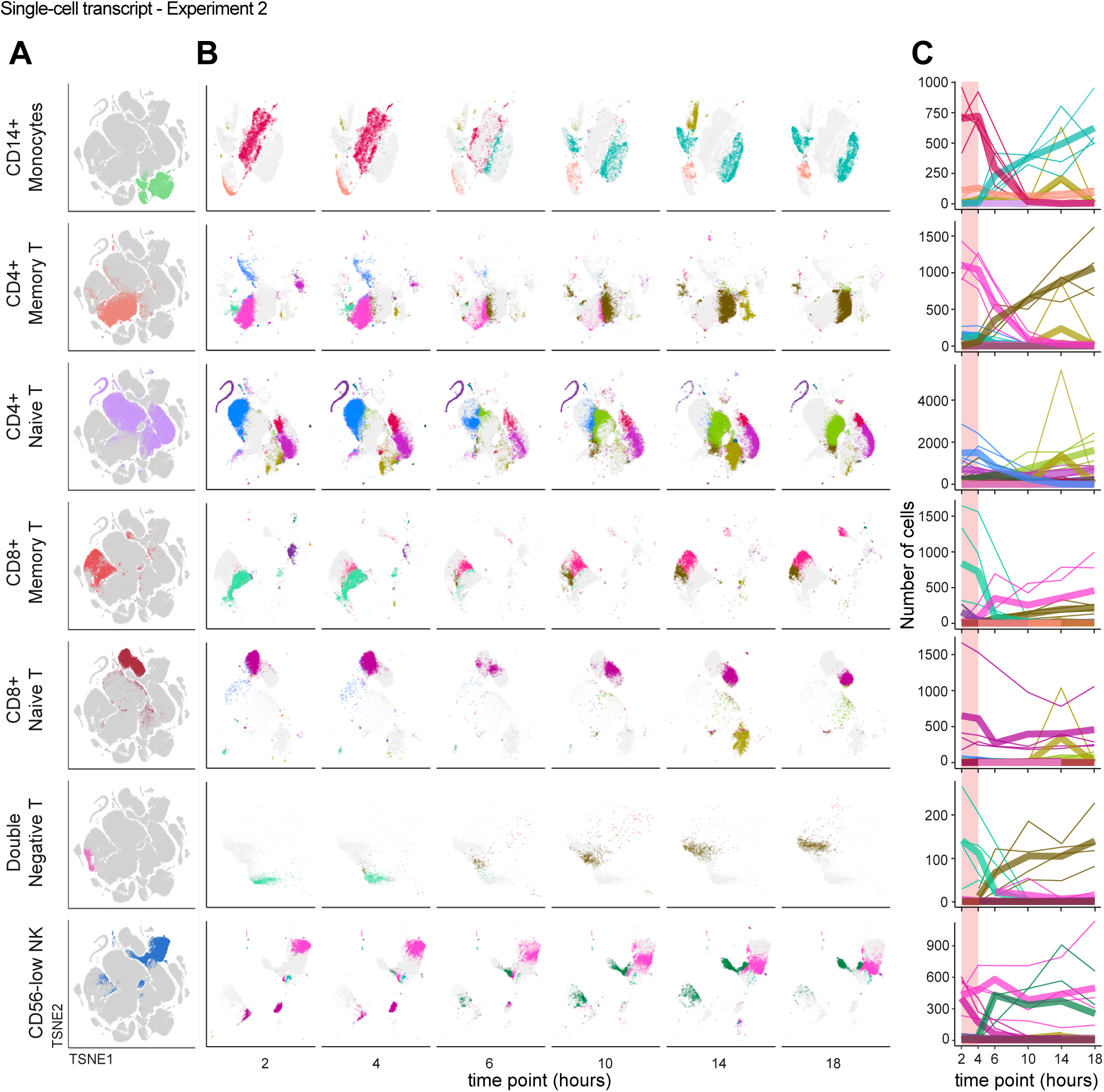
Individual cell types undergo distinct but analogous paths away from the native transcriptional state over time. Single-cell RNA-sequencing data from Experiment 2 was analyzed as in Figure 4 and the Methods. (A-B) Data were isolated based on reference-based cell type, shown on the global tSNE map (A), and displayed separately based on time point using the same tSNE coordinates (B). Colors in (A) are assigned by cell type label and are arbitrary. Each color in (B) represents a different Louvain cluster. (C) The number of cells occupying each cluster were enumerated and the counts (y-axis) plotted per cell type relative to the time after blood draw (x-axis) for each donor. Colors match the Louvain cluster colors in (B). Thicker lines represent the mean for all donors and thinner lines represent single donors. The 2-4 hour range highlighted in pink shows a period of apparent stability prior to more extreme divergence in cluster occupancy after 4 hours. See also Supplemental Figure S6 and Supplemental Tables S3, S4, and S5.

Reflecting the changes in the global gene expression profile observed through unsupervised clustering, quantification of differentially expressed genes (DEG) relative to the 2-hour time point demonstrated a trend of an increasing number of DEG across cell types (**Supplemental Figure S6** and **Supplemental Table S4**). Once again, the number of changes was muted when PBMC processing started at 4 hours but emerged following the 6-hour time point. As expected, the specific transcriptional changes exhibiting time-dependent regulation were varied across cell types, but generally reflected an elevated activation status and included many shared transcripts, such as NF_κ_B pathway genes (*NFKBIA*, *REL*, *TNFAIP3*), *CD83*, JUN family members, *HIF1A*, and *EIF1*. But coordinated regulation was also apparent, such as induction of *MAP3K8*, *IRF2BP2*, *TLE3*, and *MIR22HG* in both monocyte subsets and classical dendritic cells, and *SBDS* and *RBM38*, both RNA binding proteins, in all eight lymphocyte subsets.

Down-regulated genes exhibited similar trends. *DDX17*, *RIPOR2*, *TRAF3IP3*, *UCP2*, and *CD53* were diminished across many of the 14 cell types classified in the scRNA-seq data, while *DYNLL1*, *IFI16*, *POU2F2*, *CALM2*, *FYB1*, and *FGL2* were coordinately down-regulated in monocytes and dendritic cells in both experiments (**Supplemental Table S4**). We were particularly interested in the GIMAP (GTPase of the immunity-associated protein) family, a collection of eight sequence-related genes implicated in pro- and anti-apoptotic functions, primarily through studies in mouse T cells (Filén and Lahesmaa, 2010). *GIMAP4* and *GIMAP7* were down-regulated in nearly all cell types, with the exception of B cells and plasmacytoid dendritic cells, and *GIMAP1*, *GIMAP6*, and *GIMAP8* also exhibited decreasing expression in various cell types over the time course. In fact, *GIMAP4*, *GIMAP7*, and *GIMAP1* were among the genes that had the greatest drop in cellular percentage detection, irrespective of expression level. These data suggest survival or apoptosis may be key processes impacted by delayed PBMC processing.

To more comprehensively assess the changes occurring through the processing delay we performed pathway analysis by comparing the dominant cluster late in the time series and the dominant cluster early in the time series within a given cell type. For this analysis we selected CD14^+^ monocytes and CD4^+^ memory T cells as examples of myeloid and lymphoid cell types frequently interrogated in systems biology studies. Through this analysis, 18 of the top 22 pathways in CD14^+^ monocytes and 23 of the top 26 pathways in CD4^+^ memory T cells enriched in the late clusters included NF_κ_B and/or JUN family members (**Supplemental Table S5**), indicating the emergence of an activated phenotype within these cell types. Also of note was the elaboration of pathways in both cell types related to HMGB1, hypoxia, Th17 activation, the NFκB pathway, IL-6 signaling, and apoptosis. Other pathways of note include two AHR pathways and TREM1 signaling induced in CD14^+^ monocytes, and the mTOR and unfolded protein response in CD4^+^ memory T cells. These pathways variously relate to survival and apoptosis, inflammation, and sensing of extracellular signals that potentially confound the native *ex vivo* biology. Though we cannot determine whether this shift in cluster occupancy and gene expression program represents a direct precursor progeny relationship or whether it results from the outgrowth of a distinct rare subset of cells within each population, in either case it is clear that cells labeled as CD14^+^ monocytes or CD4^+^ memory T cells have undergone a time-dependent shift in their transcriptional profile. On the whole, the single-cell transcriptomics data demonstrate that cells are not “resting” in whole blood in the time between blood draw and PBMC processing, but rather are subject to and participating in a complex and active immune environment.

### Aligned -omics data powers hypothesis generation

A unique advantage of these data is the ability to use them in cross-modal analyses. All data sets from a given donor were acquired from the same blood draw, eliminating subclinical immunity, circadian, diet, sleep, seasonal allergy, and a host of other environmental variables as confounders for intra-donor longitudinal analysis. In addition, targeted bulk transcriptomics and single-cell transcriptomics were performed from the same PBMC isolation per time point. Therefore, these single-cell RNA-seq data are ideal for deconvoluting the bulk transcriptomics of PBMC. One striking example is the T cell costimulatory molecule ICOS ligand (encoded by *ICOSLG*). Targeted bulk transcriptomics indicates a modest log2 fold change of 1.7-2.3 across the six donors in Experiment 1 (Figure 6A). Single-cell RNA sequencing, however, identifies induction of *ICOSLG* not only in myeloid cells such as conventional dendritic cells and plasmacytoid dendritic cells but also in B cells, with the highest expression in naive B cells (Figure 6B). Another interesting example is the secreted plasminogen activator urokinase, encoded by the gene *PLAU*, which was identified in the bulk transcriptomics data for a moderate induction by the 18-hour time point of Experiment 1. Single-cell analysis indicates the majority of this induction occurs in plasmacytoid dendritic cells, with little expression in other cell types. Urokinase is localized to the extracellular plasma membrane by association with PLAUR (Ellis and Danø, 1991), which was likewise induced in bulk transcriptomics and shown to be expressed by monocytes and dendritic cells in our data, thus establishing a potential source and reservoir for potentiation of thrombolytic activity. These examples highlight the potential to explicitly decode bulk transcriptomic data sets of PBMC using our six matched longitudinal data sets.

**Figure 6.**
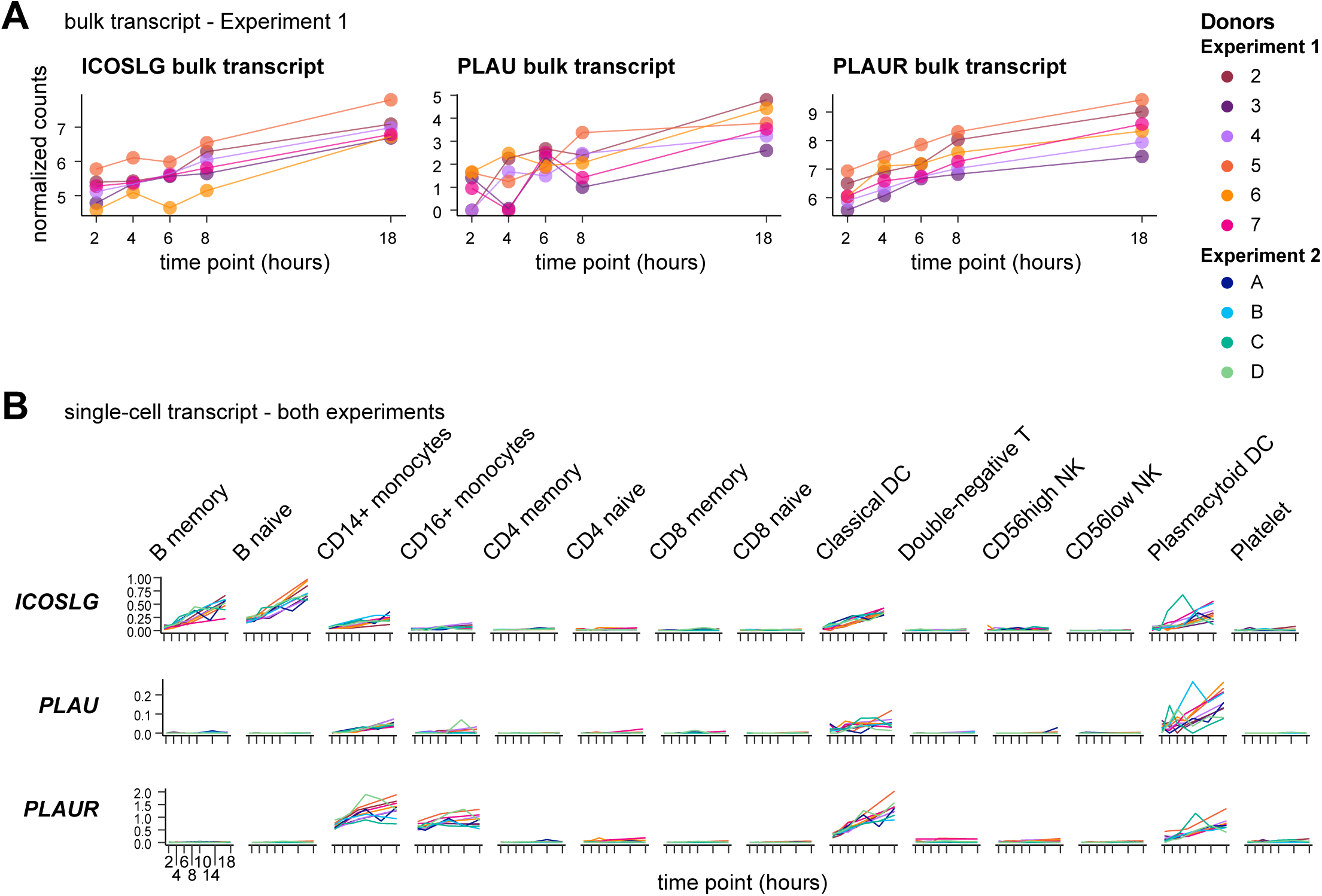
Deconvolution of bulk transcriptomics by paired single-cell RNA-seq enables identification of cell-type-specific features. (A) Transcripts from bulk transcriptomics data analyzed as in Figure 2 were selected based upon increasing expression over time and consistent dynamics across all six donors in Experiment 1. Charts show normalized counts for hours since blood draw for the individual donors, indicated by color. (B) Single-cell RNA-seq data (analyzed as in Figure 4) from all 10 donors in both experiments were queried for transcripts selected in (A) and plotted by cell type. Charts show the normalized and scaled RNA count (y-axis) for hours since blood draw (x-axis). Individual donors are indicated by color. See also Supplemental Tables S1 and S4.

More intriguing is the potential for synergy with cross-modal data. Whole blood left on the benchtop to be processed is a closed system and observations of concordant changes in a plasma protein and its corresponding transcript within a given cell type could be used to build testable hypotheses. To find such protein-transcript relationships, we filtered the list of proteins identified in Figure 3 for those having significant slopes over the entire time course, and determined the significance and direction of the slopes of the corresponding transcripts in each cell type. Though the number of anti-correlations outnumbered correlations, we focused on the proteins that were increasing over the time course and their positively correlated transcripts, as such a trend suggests *de novo* transcription, translation, and release resulting from activity occurring after blood draw. 137 proteins (11.8%) were significantly increased over the time course, of which approximately 25% were correlated with their corresponding transcript in at least one cell type and almost 11% in 3 or more cell types (**Supplemental Figure 7** and **Supplemental Table S6**). A limited number of protein-transcript pairs were increased in 9 or more of the 14 populations: PTPN1, NFKBIE, ELOA, TR, BACH1, RPS6KB1, and CD69. Conversely, the only decreased protein showing concordance with its transcript was APLP1, the strongest decreased protein in both experiments (Figure 3), and the correlation was found only with naïve B cells. Though the number of correlations overall was low, these data are consistent with the hypothesis that transcriptional activity *ex vivo* can lead to protein changes in the blood, with potential for cascading effects on bystander cells.

To consider these correlations in a more hypothesis-generating approach, we highlight two targets from this analysis. One, the TNF superfamily member LIGHT (encoded by *TNFSF14*), is a membrane and secreted ligand of the receptors HVEM, LTβR, and the soluble decoy DcR3. LIGHT potentiates T cell responses (Shaikh et al., 2001), has been reported in the setting of autoimmunity (Herro et al., 2015; Kotani et al., 2012), and is actively pursued as an anti-cancer therapy (Skeate et al., 2020). In our proteomics data, detectable LIGHT protein in the plasma fraction increased over the blood processing delay time course of both experiments (Figure 7A), resulting in a mean 2.49 fold increase at 18 hours post-blood draw relative to 2 hours, the 20^th^ most-induced protein in that comparison. Simultaneously, single-cell RNA sequencing showed the highest expression of LIGHT in CD8^+^ memory αβT cells, NK cells, and double-negative T cells (likely comprised of CD4 CD8 372 αβT and many γδT cells) (Figure 7C). One of the receptors of LIGHT, HVEM (encoded by *TNFRSF14*), was broadly expressed but the other major receptor *LTBR* was restricted primarily to the myeloid compartment. At the same time, the decoy receptor DcR3 (encoded by *TNFRSF6B*), capable of negatively regulating LIGHT (Wroblewski et al., 2003), was expressed primarily in lymphocytes, especially by double-negative T cells, which showed a trend toward increasing transcript levels concordant with increased levels of LIGHT plasma protein.

**Figure 7.**
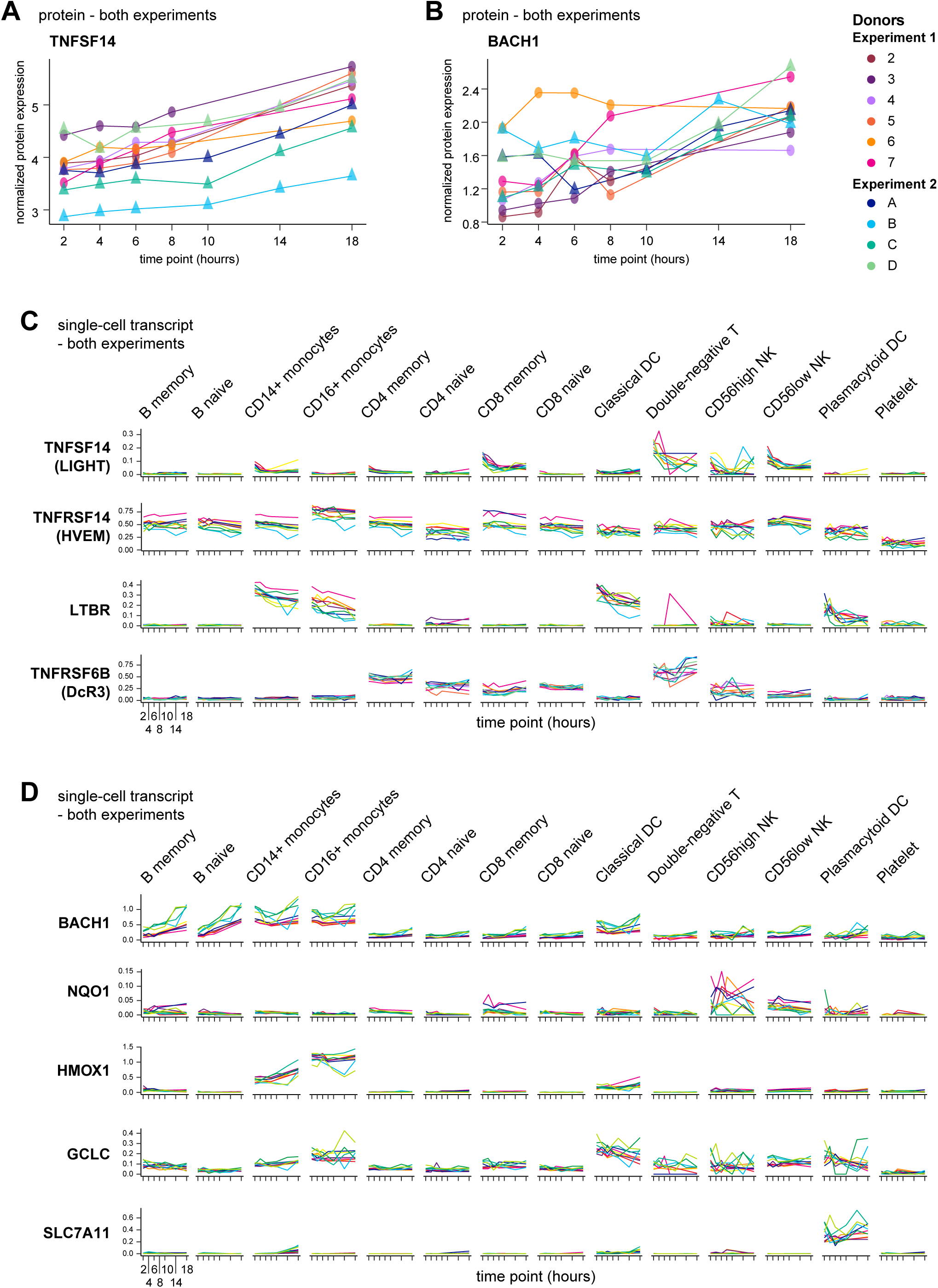
Multi-modal analysis reveals unanticipated protein-gene-cell relationships. (A-B) Plasma proteins from proteomics data analyzed as in Figure 3 were selected based upon increasing expression over time and across all 10 donors in both experiments. Normalized protein expression (y-axis) over time since blood draw (x-axis) is shown, with individual donors indicated by color. (C-D) Single-cell RNA-seq data (analyzed as in Figure 4) from all 10 donors in both experiments were queried for transcripts encoding TNFSF14 and its receptors (C) or BACH1 and a selection of its transcriptional targets related to oxidative stress (D), and plotted by cell type. Charts show the normalized and scaled RNA count (y-axis) for hours since blood draw (x-axis). Individual donors are indicated by color. See also Supplemental Figure S7 and Supplemental Table S8.

Another intriguing participant in the complex *ex vivo* blood environment is BACH1. BACH1 is a BTB/POZ transcription factor that heterodimerizes with small MAF proteins, leading to the regulation of oxidative stress gene targets bearing MAF recognition elements (Zhang et al., 2018). One such target, *HMOX1*, can be rapidly induced in monocytes by inflammatory stimuli (Yachie et al., 2003), and in fact we recognized induction and expression of *HMOX1* in CD14^+^ and CD16^+^ monocytes over the processing delay in our data (Figure 7D). BACH1 was identified in our data sets by an increase in plasma protein levels (Figure 7B) and has previously been recognized as a regulator of *HMOX1* in human monocytes (Miyazaki et al., 2010). Consistent with the literature, we found the highest levels of *BACH1* transcript were expressed by monocytes, especially early after blood draw (Figure 7D). However, whereas nearly all other populations exhibited little to no expression of *BACH1* throughout the time course, both naive and non-naive B cells demonstrated a sharp induction of *BACH1* gene expression. A role for BACH1 in B cells has not been reported aside from potential redundancy with BACH2 during B cell development (Itoh-Nakadai et al., 2014). Thus, this observation represents a potential novel aspect of B cell biology under conditions of cellular stress and highlights the potential for these aligned data sets not only to reveal putative sources of activation-induced proteins, but to suggest possible cellular networks of communication.

## Discussion

The human immune system can be biopsied through the collection of peripheral blood samples, making blood one of the most amenable human tissues for research studies. As a ‘sense and respond’ organ, the immune system tailors its activity to a wide variety of subtle environmental cues, including *ex vivo* manipulation (Kelley et al., 1987). We used a longitudinal multi-modal approach to better understand artifacts from process variability in the study of whole blood components, providing three important contributions to the study of the human immune system. First, we showed that a delay in processing venipuncture blood samples has profound consequences on immune cells. Global effects on the native biology were identified in multiple assays, in all 10 donors of the study, and in all cell types we identified. Second, our multi-modal and longitudinal data sets constitute over 10 billion measurements of immune features in a closed system. These data can be mined for correlative gene-gene, gene-protein, ligand-receptor, and pathway interactions from which testable hypotheses can be derived. Furthermore, as these activities relate to potentially confounding pre-analytical artifacts, these data may enable a qualification of results from other studies that are subject to variable or sub-optimal processing conditions. Third, the 2-hour time points provide a large, high-quality, multi-modal data set of 10 human donors. These 2-hour data are temporally close to the *in vivo* biology, consisting of flow cytometry for major PBMC populations, single-cell RNA-sequencing of ∼8000 quality cells per donor, targeted plasma proteomics of >1000 targets, and targeted bulk transcript analysis of >600 targets on some donors. Thus, this resource provides various opportunities to investigate a multi-modal dataset derived from the components of freshly isolated and *ex vivo* aged human blood.

While the data from typical flow cytometry implied that commonly used cell surface proteins and the cells themselves are stable, both the targeted as well as whole transcriptome studies confirm large-scale changes in gene regulation. After an 18-hour delay in blood processing, transcriptional states in all cell types were fundamentally altered. Given the length of time, this might be expected. But more concerning are the changes occurring after just 4 and 6 hours post-draw, as evidenced by the changes in cellular state apparent through single-cell RNA-sequencing. Our findings implicate an up-regulation in inflammation and stress-activation (*NFKBIA*, *HIF1A*, *MAP3K8*), perhaps in response to deteriorating environmental conditions (Paardekooper et al., 2018; Rius et al., 2008). Under these conditions, the appearance of new clusters in tSNE visualization of gene expression data likely results from a large number of cells altering their gene expression profiles *ex vivo* rather than outgrowth of more rare pre-existing cells faithfully representing an *in vivo* activation state. If true, this fact underscores the challenges of using tSNE/UMAP classification of the data as an analysis mode, as cluster occupancy can be confounded by blood processing delay. The preferred alternative is to identify cellular identity on a per-cell basis first, and then compare the transcriptomic state of equivalent cell types based on the experimental variable (*e.g.* disease state); in this case the sample processing metadata, such as blood processing delay, must be carefully tracked and corrected for in the analytical model.

As predicted from our gene expression data, the plasma proteome exhibited dramatic changes in concert with the PBMC transcriptomics data. Correlation analysis targeted at identifying induction of transcripts coding for proteins that were coordinately increased revealed a number of interesting relationships, including some proteins known to be highly relevant to immunity (PTPN1, NFKBIE, TR, BACH1, CD69). However, it was apparent that anti-correlations were more prevalent than correlations, perhaps reflecting negative feedback in the presence of elevated protein signal, positive feed-forward when signal becomes limiting, or unlinked transcriptional regulation in the context of pre-formed protein. This result underscores the importance of studying protein pro-forms and post-translational regulation in order to establish explicit relationships between transcripts and their bioactive protein products.

Efforts to understand the effect of processing delay on various plasma and serum analytes have existed for some time (Ignjatovic et al., 2019; Ono et al., 1981), though typically limited to individual proteins or metabolites of known clinical interest. We have generated a unique public human proteomic data set, consisting of over 62,000 data points linked to PBMC flow cytometry and transcriptomics. The clearest signal in the targeted proteomics relates to platelet biology. This includes CD40L, PDGFβ, ANGPT1, STK4, and others, supportive of a hypothesis of early and ongoing platelet activation which likely results in either direct and indirect protein release or increased platelet contamination in the plasma isolation. Blood for the proteomics studies was drawn into K2-EDTA tubes, the recommended anticoagulant for the assay and consistent with common practice. EDTA has previously been reported to elicit aggregation of platelets *in vitro* (Pegels et al., 1982). Of note, platelet aggregation in whole plasma is also not inhibited by heparin (Saba et al., 1984), such that the platelet response to blood draw or hemostasis may elicit or potentiate divergence from native biology. Therefore, our data suggests that some of the most severe changes in plasma proteomics result from the basic process of obtaining the sample. This activity has the potential for cascading effects on PBMC response, including by binding to and activating various leukocytes, serving as a physical mark of inflammation, and providing key signals such as CD40 ligand, CCL5, and CXCL4 (Gaertner and Massberg, 2019). These changes are exacerbated by time delay, thus highlighting the need for rapid and consistent processing.

In order to mitigate these confounding effects, the optimal solution is to process the samples consistently and with as little delay as possible. Previous studies on bulk PBMC have suggested that storing samples on ice may mitigate the effects of delayed blood processing (Goods et al., 2018), and so to test this hypothesis we kept some of the whole blood samples at 4°C in a parallel experiment (data not shown). We found this mitigation effort to have mixed results, with an unexpected decrease in PBMC yield and hemolysis occurring in some samples. Therefore, our data indicates processing within 4 hours should become standard practice for assays using human blood components, particularly those employing deep transcriptional or proteomic profiling. This may not be possible in all cases. In these situations, our data provide a comprehensive set of landmarks to qualify the analysis for processing delay artifacts.

Embedded within the larger resource provided here, the 2-hour cytometry, proteomic, and transcriptomic data sets from 10 donors is a substantial resource in its own right, with minimal and controlled processing delay. The available public cytometry and single-cell RNA-seq data on human PBMC continue to grow in size and complexity, but to our knowledge no study integrates these two modalities with deep proteomics at this scale. Furthermore, though blood processing delay results in immune processes that may misrepresent or obscure the true biology being studied, we suspect that in many cases this ‘artifactual’ biology echoes physiologic processes. This potential was highlighted by the broad down-regulation of multiple GIMAP family genes in many cell types. As these are known to relate to apoptosis in T cells (Filén and Lahesmaa, 2010), this observation is interesting. However, our data also opens the possibility of a similar role in myeloid cells, for which there is little existing data (Hellquist et al., 2007; Krücken et al., 1997). Therefore, the multi-modal data set is a substrate for generating testable hypotheses, so long as the limitations of an *ex vivo* artificial system are fully accepted. In particular, blood flow has ceased and cells may aggregate by settling, there is no regulation of blood gases, and temperature is well below physiologic range. When approached with an understanding of the experimental conditions employed, we believe these data can be used productively, for example in the study of transcriptional co-regulation, correlations among proteins and between proteins and genes, as well as novel associations of genes with cell types.

Recently, Massoni-Badosa *et al*. published a complementary effort on the effect of blood processing delay (Massoni-Badosa et al., 2020), with differences in the assays, cell numbers, and analyses performed. Our overall conclusions align with theirs, in the clear risks to data interpretation from processing delay and non-uniformity, though we did not recognize a global or cell type-specific reduction in transcript levels (**Supplemental Figure S4**). Nonetheless, the practical realities of obtaining clinical samples will ultimately dictate how and when they are processed, and the data generated should be analyzed with an appreciation for the unknown and/or variable amount of delay in processing. While this naturally applies to myriad ongoing and historical clinical studies, it is also keenly relevant to public health imperatives such as the SARS-CoV2 pandemic, for which process standardization may be outweighed by the benefit of sample acquisition. Accounting for such practicalities in a data-driven manner will provide broad and on-going benefit for improving the caliber of putative therapeutic targets, by lowering confidence in those subject to *ex vivo* regulation and thereby increasing confidence in the others. We hope these data will also spur a renewed focus on process standardization and optimization to ultimately improve data quality and physiologic relevance at the point of data generation. With over 1 billion data points across several -omic platforms, our resource offers a rich data repository upon which to qualify existing data, benchmark and train subsequent studies, and derive novel hypotheses.

## Supporting information

Savage Figure S1-PBMC metrics, flow cytometry gating

Savage Figure S2-Flow cytometry frequencies

Savage Figure S3-Nanostring bulk transcriptomics PCA

Savage Figure S4-Single-cell RNAseq, technical metrics

Savage Figure S5-Single-cell RNAseq, tSNE by time

Savage Figure S6-Single-cell RNAseq, DEG

Savage Figure S7-Protein-transcript correlations

Savage Figure S8-technical variance

Savage Table S1-Nanostring MEM complete

Savage Table S2-Olink complete

Savage Table S3-scRNAseq_QCstats_clusters_coords

Savage Table S4-scRNAseq-Significant_DEG

Savage Table S5-scRNAseq-pathways

Savage Table S6-Protein-transcript correlations

Savage Table S7-Donor information

Savage Table S8-scRNAseq_Significant_DEG

## Acknowledgements

We are grateful to the individuals that provided biological material for this study. We appreciate the support of David Skibinski and Deric Khuat at Benaroya Research Institute. We thank Olink Proteomics for providing complimentary proteomic studies. We are especially thankful to all the members of the Allen Institute for Immunology and the facilities and operations teams at the Allen Institute who helped establish the productive environment in which this work was performed. We are grateful for the leadership and support of Allan Jones, President and CEO of the Allen Institute, and to the Allen Institute for Immunology for funding this study. The authors also wish to thank the Allen Institute founder, Paul G. Allen, for his vision, encouragement, and support.

## Author Contributions

T.F.B., M.V.G., A.K.S., and H.B. conceived the study. M.V.G., A.K.S., H.B., E.S., T.C., and P.J.S. designed the experiments. A.K.S., A.T.H., K.H., N.K., K.C.B., T.S., and M.V.G. prepared the PBMC and plasma from whole blood. A.K.S. and A.T.H. performed the flow cytometry. A.K.S. and K.H. analyzed the flow cytometry data. E.S., C.L., and P.C.G. performed the single-cell RNA-sequencing. M.V.G., T.C., R.G., M.C., X.L., H.B., Z.T., and J.G. analyzed the single-cell RNA-sequencing data. M.V.G., M.C., and X.L. analyzed the bulk transcriptomics data and proteomics data. M.V.G. and A.K.S. selected candidates for cross-modal examination. E.J. built the web-accessible data browser. A.B. managed computational resources. J.H.B. provided human samples. T.F.B., P.J.S., T.R.T., X.L., E.M.C, and P.M. provided direction and oversight. A.K.S. and T.C. wrote the manuscript, and all authors provided edits and comments to the manuscript.

## Supplemental tables

**Supplemental Table S1, Related to** Figure 2, Figure 6**, and Supplemental Figure S3 – Nanostring bulk transcriptomics**

**Supplemental Table S2, Related to** Figure 3**, Supplemental Figure S7, and Supplemental Table S8 – Olink proteomics**

**Supplemental Table S3, Related to** Figure 4 **and Supplemental Figure S4 – single-cell RNA sequencing, per-cell metrics and metadata**

**Supplemental Table S4, Related to** Figure 5, Figure 6**, and Supplemental Figure S5 – single-cell RNA sequencing, differentially expressed genes relative to the 2-hour time point, by cell type**

**Supplemental Table S5, Related to** Figure 5 **– single-cell RNA sequencing, pathway analysis**

**Supplemental Table S6, Related to Supplemental Figure S7, and Supplemental Tables S2 and S4 – Protein-transcript correlations**

**Supplemental Table S7 – donor information**

**Supplemental Table S8, Related to** Figure 5 **and Supplemental Figure S5 – single-cell RNA sequencing, differentially expressed genes by cell type classification**

## Supplemental figure and legends

**Supplemental Figure S1, Related to** Figure 1 **– PBMC yield and viability, flow cytometry gating**

(A-B) PBMC were isolated from whole blood by ficoll separation at the indicated time points after blood draw and assessed for yield (A) and viability (B).

(C-D) 5×10^6^ PBMC isolated as in panels A-B and frozen in liquid nitrogen were thawed and assessed for yield (C) and viability (D) prior to downstream assays. “% recovery” was calculated as the recovery of live PBMC divided by 5×10^6^.

(E-F) Flow cytometry gating schemes for Experiment 1 (E) and Experiment 2 (F). Orange labels indicate gates used to determine population frequencies. The gate labeled “cleanup” in Experiment 2 was used to remove dye aggregates.

**Supplemental Figure S2, Related to** Figure 1 **– Flow cytometry frequencies**

Flow cytometry data were gated by traditional methods for major PBMC populations, as in Supplemental Figure 1. Because of technical artifacts, the 6-hour time points of donors A and B were excluded from analysis.

(A-B) The percent change in frequency was calculated for each population relative to the 2-hour PBMC processing time point and population medians were calculated. Data are displayed as a heat map for Experiment 1 (A) or Experiment 2 (B).

(C-D) Population frequencies are shown per donor as a percent of live cells in Experiment 1 (C) or percent of CD45^+^ cells in Experiment 2 (D).

**Supplemental Figure S3, Related to** Figure 2 **and Supplemental Table S1 – Nanostring bulk transcriptomics principle components analysis**

PBMC in Experiment 1 were prepared from whole blood at various times after blood draw and assayed by targeted transcriptomics in bulk using Nanostring nCounter. The normalized counts were analyzed by principle components analysis and colored by donor (A) or by time point (B).

**Supplemental Figure S4, Related to** Figure 4 **and Supplemental Table S3 – Single-cell RNA sequencing, technical metrics**

Single-cell RNA-sequencing technical metrics were compiled for Experiment 1 (A-D) and Experiment 2 (E-H) and categorized by cell type, as assigned by Seurat-based reference alignment (see Methods for details).

The number of quality singlets (A, E), mean mitochondrial DNA (B, F), features (C, G), and UMI (D, H) were quantified per cell type and colored by donor.

**Supplemental Figure S5, Related to** Figure 4 **and Supplemental Table S3 – Single-cell RNA sequencing, tSNE individually by time**

The single-cell RNA-sequencing normalized gene expression matrices from each sample were identified for donor and time point and aggregated per experiment. The multidimensional data was displayed in two dimensions using tSNE, split out by time post-blood draw for Experiment 1 (A) and Experiment 2 (B), and colored by Louvain cluster. Additional details can be found in the Methods.

**Supplemental Figure S6, Related to** Figure 5 **and Supplemental Tables S3, S4, and S5 – Single-cell RNA sequencing, differentially expressed genes**

Differential expression in genes at each time point relative to 2 hours post-draw are represented as log2 fold change (x-axis) vs. log10 adjusted p-value (y-axis) (see the Methods for details). Blue dashed lines indicate log2 fold change of |1.1| and red dashed lines indicate adjusted p-values of 0.05 and 0.01. For visualization purposes, adjusted p-values at zero are set to the non-zero minimum of all other adjusted p-values and divided by two. Cell types not represented did not have differentially expressed genes (after p-value adjustment).

(A-C) Differential expression in healthy (A) and SLE (B) donors of Experiment 1 and healthy donors of Experiment 2 (C). The legend in panel (A) also applies to panel (B).

**Supplemental Figure S7, Related to** Figure 7**, and Supplemental Tables S2, S4, and S6 – Protein-transcript correlations**

Proteins identified in Figure 3 were filtered for significance and directionality across the entire time course and the corresponding transcripts were similarly assessed by cell type.

(A) Proteins of increasing and decreasing abundance were scored for correlation (black) with the corresponding transcript in a least one cell type. Protein-transcript pairs that were not correlated (gray) could be anti-correlated, not significant, or have missing transcript data (for example, from drop-outs in the single-cell RNA-sequencing data).

(B) To assess whether protein-transcript correlations were common to many cell types or typically restricted to a limited number of cell types, increasing (orange) and decreasing (purple) proteins were scored for correlation (solid color) or anti-correlation (hashed color) with their corresponding transcript in each cell type and the count of each correlation was tallied across the cell types. Because proteins are scored if they are correlated or anti-correlated in at least one of fourteen populations, a single protein could be scored as both.

(C-D) To assess whether specific cell types had a propensity or bias in their correlations, correlated (solid color), anti-correlated (hashed color), and not correlated proteins (gray) were tallied by cell type for proteins increased (C, orange) and decreased (D, purple) over the time course.

**Supplemental Figure S8, Related to Supplemental Tables S2 and S4 – Analysis of technical variance in proteomics and single-cell RNA-sequencing assays**

(A-B) To assess the technical variance of the plasma protein assay, plasma samples from six donors from both 2 hours and 6 hours post-blood draw were compared across three different assay plates. The data are shown as MA plots (A) and the distributions of fold changes (B) in plate-to-plate comparisons. The red dash lines indicate the log2 fold change cutoff (0.585) used in the study.

(C-D) To assess the technical variance of the single-cell RNA-sequencing assay, replicate aliquots of PBMC from a non-study sample were assayed in three different wells (see Methods/Single Cell Transcriptomics). Average transcript intensities were calculated from cells with non-zero counts and fold changes between replicates were calculated. The data are shown as MA plots (A) and the distributions of fold changes (B) in well-to-well comparisons. The red dash lines indicate the log2 fold change cutoff (1.1) used in the study.

## Methods

### Resource availability

#### Lead Contact

Further information and requests for resources and reagents should be directed to and will be fulfilled by the Lead Contact, Adam Savage.

#### Materials Availability

No materials were generated for this study.

#### Data and Code Availability

Flow cytometry .fcs data files are available on request. Nanostring bulk transcriptomics and Olink proteomics data are provided in **Supplemental Tables S1 and S2**, respectively. All single-cell RNA-sequencing data have been deposited at GEO (GSE156989). Transcriptomics and proteomics data are available for exploration at http://bloodprocessingdelay.allenimmunology.org.

### Methods Details

#### Donors and sample handling

Blood samples were obtained from healthy (no diagnosis of disease, donors 2-4 and A-D) and non-matched donors with systemic lupus erythematosus (SLE, donors 5-7) (**Supplemental Table S7**), from the Benaroya Research Institute (Seattle, WA) or Bloodworks Northwest (Seattle, WA) through protocols approved by the relevant institutional review boards. Blood was drawn into BD NaHeparin vacutainer tubes (for PBMC; BD #367874) or K2EDTA vacutainer tubes (for plasma; BD #367863) and, upon arrival at the processing lab, all NaHeparin tubes for each donor were pooled into a sterile plastic receptacle to establish one common pool for all time points and stored at room temperature for the duration. PBMC isolation and plasma processing was started at 2, 4, 6, 8, and 18 hours post-draw (Experiment 1) or 2, 4, 6, 10, 14, and 18 hours post-draw (Experiment 2). For Experiment 1, each donor sample was processed by a single operator on a separate day, and thawed PBMC were assayed by cytometry and single-cell RNA-sequencing in three batches (donor 2, donors 3-4, and donors 5-7). For Experiment 2, all four donors were processed on the same day by a team of operators, and thawed PBMC were assayed by cytometry and single-cell RNA-sequencing in one batch. Due to a PBMC processing error in Experiment 2, flow cytometry and single-cell RNA-sequencing was not available for the 6-hour time point from donors A and B. This resulted in 30 samples of bulk transcriptomics (Experiment 1 only), 52 samples of flow cytometry and single-cell RNA-sequencing, and 54 samples of plasma proteomics data.

For PBMC isolation, at each time point the pool of blood was gently swirled until fully mixed, about 30 times, and a volume of blood was removed and combined with an equivalent volume of room temperature PBS (ThermoFisher #14190235). PBMC were isolated using one or more Leucosep tubes (Greiner Bio-One #227290) loaded with 15 mL of Ficoll Premium (GE Healthcare #17-5442-03) to which a 3 mL cushion of PBS had been slowly added on top of the Leucosep barrier. The 24-30 mL diluted whole blood was slowly added to the tube and spun at 1000xg for 10 minutes at 20°C with no brake. PBMC were recovered from the Leucosep tube by quickly pouring all volume above the barrier into a sterile 50 mL conical tube. 15 mL cold PBS+0.2% BSA (Sigma #A9576; “PBS+BSA”) was added and the cells were pelleted at 400xg for 5-10 minutes at 4-10°C. The supernatant was quickly decanted, the pellet dispersed by flicking the tube, and the cells washed with 25-50 mL cold PBS+BSA. Cell pellets were combined, if applicable, the cells were pelleted as before, supernatant quickly decanted, and residual volume was carefully aspirated. The PBMC were resuspended in 1 mL cold PBS+BSA per 15 mL whole blood processed and counted with a ViCell (Beckman Coulter) using VersaLyse reagent (Beckman Coulter #A09777) or with a Cellometer Spectrum (Nexcelom) using Acridine Orange/Propidium Iodide solution. PBMC were cryopreserved in Cryostor10 (StemCell Technologies #07930) or 90% FBS (ThermoFisher #10438026) / 10% DMSO (Fisher Scientific #D12345) at 5×10^6^ cells/mL by slow freezing in a Coolcell LX (VWR #75779-720) overnight in a −80°C freezer followed by transfer to liquid nitrogen.

For plasma isolation, the K2-EDTA source tube was gently inverted 10 times and the appropriate volume of whole blood was extracted using an 18-gauge needle and syringe, and transferred to a similar plastic tube with no additives (Greiner Bio-One #456085). The blood was centrifuged at 2000xg for 15 minutes at 20°C with a brake of 1, and 80-90% of the plasma supernatant was removed by careful pipetting for immediate freezing at −80°C. Plasma was assayed after the first freeze/thaw.

#### Flow cytometry

PBMCs were removed from liquid nitrogen storage and immediately thawed in a 37°C water bath. Cells were diluted dropwise with 37°C AIM V media (Thermo Fisher Scientific #12055091) up to a final volume of 10 mL. A single wash was performed in 10 mL of PBS+BSA, pelleting cells at 400xg for 5-10 minutes at 4-10°C. PBMCs were resuspended 2 mL in PBS+BSA, counted on a ViCell or Cellometer Spectrum, as above, and 1×10^6^ cells were incubated sequentially or together with Human Trustain FcX (BioLegend #422302) and Fixable Viability Stain 510 (BD #564406), on ice and according to manufacturer’s instructions. Cells were washed with PBS+BSA and stained with a master mix cocktail of antibodies on ice for 25-30 minutes. Experiment 1 and Experiment 2 were assayed by flow cytometry using a 12-target panel and a 24-target panel, respectively (Key Resources Table). Cells were washed with PBS+BSA and fixed with 4% paraformaldehyde (Electron Microscopy Sciences #15713) for 12-15 minutes at room temperature. Cells were washed with PBS+BSA, resuspended in PBS+BSA, and collected on a BD Symphony cytometer. After compensation, cytometry data was pre-processed to remove unrepresentative events due to instrument fluidics variability (time gating), to exclude doublets (by FSC-H and FSC-W), and to exclude cells exhibiting membrane permeability (live/dead gating) prior to quantification using BD FlowJo software. Gating examples are provided in **Supplemental Figure 1E-F**. The percent change in frequency was calculated per cell type to quantify the observed cell frequency changes along the variable of time, along with the corresponding change in median frequencies across all samples.

#### Bulk transcriptomics

For Experiment 1, 2×10^5^ fresh PBMC were pelleted, resuspended in Qiagen buffer RLT (Qiagen #79216), and quick frozen immediately at −80°C for assay on the Nanostring nCounter platform, performed as a fee-for-service by Nanostring using their standard protocols for the nCounter Gene Expression – Hs Immunology v2 CodeSet assay.

#### Proteomics

Plasma samples were assayed using the Olink Proximity Extension assay, run on the Fluidigm Biomark system. They were submitted to Olink Boston (Experiment 1) or Olink Sweden (Experiment 2). Patient samples were distributed evenly across plates, and all timepoints per patient were run on the same plate, with randomized well locations. Samples were assayed using the Olink Discovery Assay which encompasses all proteins across 13 panels (Cardiometabolic (V.3603), Cardiovascular II (V.5006), Cardiovascular III (V.6113), Cell Regulation (V.3701), Development (V.3512), Immune Response (V.3202), Inflammation (V.3021), Metabolism (V.3402), Neuro Exploratory (V.3901), Neurology (V.8012), Oncology II (V.7004), Oncology III (V.4001), Organ Damage (V.3311)). Data were normalized by Olink using an Olink-provided plasma plate bridging control, three positive controls, and three background controls. Samples reported below the stated Limit of Detection were removed from our analysis.

#### Single Cell Transcriptomics

PBMC were thawed as described in the Methods for flow cytometry and the same vial of cells was used for flow cytometry and single-cell RNA-sequencing. Single-cell RNA-seq libraries were generated using the 10x Genomics Chromium 3’ Single Cell Gene Expression assay (#1000075 or #1000121) and Chromium Controller Instrument according to the manufacturer’s published protocol. 16,000 cells from each PBMC sample were loaded into a separate Chromium Single Cell Chip B (10x Genomics #1000073) well targeting a recovery of 10,000 cells. Gel Beads-in-emulsion (GEMs) were then generated using the 10x Chromium Controller. The resulting GEM generation products were then transferred to strip tubes and reverse transcribed on a C1000 Touch Thermal Cycler programmed at 53°C for 45 minutes, 85°C for 5 minutes, and then held at 4°C. Following the reverse transcription incubation, GEMs were broken and the pooled single-stranded cDNA fractions were recovered using Silane magnetic beads (Dynabeads MyOne SILANE #37002D). Purified barcoded, full-length cDNA was then amplified with a C1000 Touch Thermal Cycler programmed at 98°C for 3 minutes, 11 cycles of (98°C for 15 seconds, 63°C for 20 seconds, 72°C for 1 minute), 72°C for 1 minute, and then held at 4°C. Amplified cDNA was purified using SPRIselect magnetic beads (Beckman Coulter #22667) and a 1:10 dilution of the resulting cDNA was run on a Bioanalyzer High Sensitivity DNA chip (Agilent Technologies #5067-4626) to assess cDNA quality and yield. A quarter of the cDNA sample (10 ul) was used as input for library preparation. Amplified cDNA was fragmented, end-repaired, and A-tailed in a single incubation protocol on a C1000 Touch Thermal Cycler programmed at 4°C start, 32°C for 5 minutes, 65°C for 30 minutes, and then held at 4°C. Fragmented and A-tailed cDNA was purified by performing a dual-sided size-selection using SPRIselect magnetic beads (Beckman Coulter #22667). A partial TruSeq Read 2 primer sequence was then ligated to the fragmented and A-tailed end of cDNA molecules via an incubation of 20°C for 15 minutes on a C1000 Touch Thermal Cycler. The ligation reaction was cleaned using SPRIselect magnetic beads (Beckman Coulter #22667). PCR was performed to amplify the library and add the P5 and indexed P7 ends (10x Genomics #1000084), using a C1000 Touch Thermal Cycler programmed at 98°C for 45 seconds, 13 cycles of (98°C for 20 seconds, 54°C for 30 seconds, 72°C for 20 seconds), 72°C for 1 minute, and then held at 4°C. PCR products were purified by performing a dual-sided size-selection using SPRIselect magnetic beads (Beckman Coulter #22667) to produce final, sequencing-ready libraries. Final libraries were quantified using Picogreen and their quality was assessed via capillary electrophoresis using the Agilent Fragment Analyzer HS DNA fragment kit and/or Agilent Bioanalyzer High Sensitivity chips. Libraries were sequenced on the Illumina NovaSeq platform using S4 flow cells. Read lengths were 28bp read1, 8bp i7 index read, 91bp read2.

### Computational Analysis

#### Nanostring model

The Nanostring platform was run on all timepoints from Experiment 1 (Subjects 2-7, Timepoints 2, 4, 6, 8, 18 hours). The data were normalized to Nanostring-provided control genes using the NanoStringNorm package (Waggott et al., 2012) and log-transformed (base 2). Genes with zero expression in more than 20% of samples were removed from further analysis. Generalized linear mixed effect models (GLMEM) were fit to the expression data of individual genes, controlling for the fixed effects of sex (male vs. female) and binary processing timepoints (4, 6, 8, 18 hours vs. 2-hour baseline), and subject-level random effects to account for individuals’ inherent variability, using Gaussian family distribution. All the models were built in the R library glmmTWB version 1.0.1. The log2 fold change from the baseline was evaluated by the corresponding time-related effect estimates. Multiple tests correction was applied to the p-values to control the false discovery rate (FDR) using Benjamini & Hochberg procedure (Benjamini and Hochberg, 1995).

### Proteomics

#### Proteomics Model

Plasma from Experiment 1 and Experiment 2 were submitted to Olink (Uppsala, Sweden) in one batch per experiment. All 2**-**hour samples from Experiment 1 were included in both batches as bridging controls. Proteins below the reported Limit of Detection or that exhibited more than 20% missingness or ranked in the lowest 20% among all proteins in expression (as defined by the sum of the corresponding normalized expression values) were disregarded from further evaluation. As recommended by Olink, expression values for each protein in Experiment 2 were subtracted by the corresponding protein-specific normalization factor, which was the median of the corresponding pair-wise expression difference as measured on the bridging controls between the two experiments.

A protein-specific GLMEM was fit to each normalized protein, controlling for the fixed effects of sex (male vs. female), age (35-55 years old vs. 25-35 years old), disease status (SLE vs. healthy), study id (Experiment 2 vs. Experiment 1) and continuous processing delay (time point), and accounting for an individual’s inherent variability through subject-level random effects. To capture the study specific non-linear temporal effects, the cubic, quadratic or linear effect of processing time and its interaction with study id were evaluated in a step-down approach using a likelihood ratio test. The dispersion was modeled as a function of plate, disease status and study id to allow non-constant variability. All the models were built in the R library glmmTWB version 1.0.1. We reported the average log2 fold-change in proteomic expression between later processing time points and the first time point (2-hour) along with the corresponding two-sided 95% CIs. Wald tests were conducted to assess significance of the change estimates and associated p-values. When a significant difference was found in the change estimates between the two studies due to experimental conditions unaccounted for, we reported study-specific change estimates. Multiple tests correction was applied to the p-values to control the FDR using Benjamini & Hochberg procedure (Benjamini and Hochberg, 1995).

#### Proteomics Technical Precision

Plasma samples from the six donors in Experiment 1 from the 2-hour and 6-hour post-blood draw time points were analyzed by Olink on three separate plates to determine the technical variance of the assay. The obtained abundance of each protein was corrected for plate effects by aligning the corresponding medians. The average and the log2 fold change of protein abundance were calculated between any two plates and summarized in **Supplemental Figure 8**.

### Single-Cell Transcriptomics

#### scRNA-seq Pre-processing

Binary Base Call (BCL) files were demultiplexed using the mkfastq function in the 10x Cell Ranger software (version 3.1.0), producing fastq files. Fastq files were then checked for quality (FastQC version 0.11.3) and run through the 10x Cell Ranger alignment function (cell ranger count) against the human reference annotation (Ensembl GRCh38). Mapping was performed using default parameters. Additional information about Cell Ranger can be found at https://support.10xgenomics.com/single-cell-gene-expression/software. Upon completion, Cell Ranger produced an output directory per file that contains the following: bam file (binary alignment file), HDF5 file (Hierarchical Data Format) with all reads, HDF file containing just the filtered reads, summary report (html and csv), cloupe.cloupe (a file for the 10x Loupe visual browser). HDF5 files (filtered) were then uploaded into the R statistical programming language (version 3.6.0) using the Seurat package (version 3.0) where normalization, scaling, integration and reference-based label transfer was performed for cell typing/classification.

#### scRNA-seq Cell Classification

Individual HDF5 files were loaded into the R statistical programming language (version 3.6.0) using Bioconductor (version 3.1.0) and the Seurat package (version 3.1.5). For simplicity, sample names were captured as a list in R and iteratively processed within a loop (refer to https://satijalab.org/seurat/ for more information). Within the loop, samples were normalized with the NormalizeData function followed by the FindVariableFeatures function with parameters: vst selection method and 2000 features. Label transfer was performed using previously published procedures (Stuart et al., 2019). Data was labeled with the Seurat reference dataset and checked for expected DEG by cell type (**Supplemental Table S8**). Labeling included the FindTransferAnchors and TransferData functions performed in the Seurat package. Labeling was performed in specified sequence where information acquired from the previous labeling was accumulated then set as anchors for the next sample, looping by time point across patients. The time point to time point variation first observed within the Nanostring data was used as the basis for this labeling strategy.

After label transfer was complete, we calculated read depth, mitochondrial percentage, and number of UMIs per sample. Despite normalization being performed during the label transfer process, the raw counts were stored with cell labels. After labeling and metrics were recorded, samples were merged together in Seurat using the merge function. The merged data structure was normalized (using NormalizeData and FindVariableFeatures functions) and then saved as an RDS for further analysis. For confirmation of cell types, we used the FindMarkers function (Seurat) and the MAST package to identify DEGs for each identified cell type against all other cell types. Unsupervised cell clustering was also performed separately for Experiment 1 and Experiment 2 using the Louvain / KNN-based method in the Seurat package.

#### scRNA-seq Differential Gene Expression Analysis

Differential expression analysis from later timepoints compared to two hours was conducted on a per reference-based cell type basis and for selected timepoints on a per Louvain-cluster basis using the MAST package (https://pubmed.ncbi.nlm.nih.gov/26653891/) and the FindMarkers function from the Seurat package (version 3.1.5). Because they were sequenced in different batches, differential expression was run separately for Experiment 1 Healthy Donors, Experiment 1 SLE Donors, and Experiment 2 Healthy Donors.

#### scRNA-seq Pathway Analysis

Pathway analysis was performed on single-cell transcriptomics data from Experiment 2. Individual cells were labeled by cell type and clustered as indicated in Methods/Single-Cell Transcriptomics/Cell Classification. As an example of pathway analysis, two clusters exhibiting extreme and inverse time-dependent bias (e.g. present in a cell type at 2 hours but not present for the same cell type at 18 hours, and vice versa) were selected and the corresponding data was submitted to Ingenuity Pathway Analysis (Qiagen, release: Summer 2020) using log2 fold change values and adjusted p-values calculated by differential gene expression analysis. The clusters selected for analysis were (colors as shown in Figure 5B): magenta (“early”) and teal (“late”) from CD14+ Monocytes and, separately, pink (“early”) and brown (“late”) from CD4+ memory T cells.

#### scRNA-seq Technical Precision

Three replicate aliquots of a non-study PBMC sample (Bloodworks Northwest, Seattle, WA) were analyzed in three wells on a Chromium Single Cell Chip B to study the precision of the technology. The average transcript intensity was calculated for each well from cells detected in the well and having nonzero counts. The obtained transcript intensity was corrected for well effects by aligning the corresponding medians of all transcripts. The average and the log2 fold change of transcript intensity were calculated between any two wells and summarized in **Supplemental Figure 8**.

#### Protein-transcript correlations

A total of 233 plasma proteins showed significant changes in any time point, of which 184 were unambiguously mapped to a single gene and detected by scRNA. The changes of these 184 proteins were then modeled as linear functions of time (linear mixed models, fixed effects: intercept and slope, random effects: intercept and slope). A total of 159 proteins showed significant slope for time (p < 0.05 after multi-testing correction). Afterwards the corresponding RNA expressions of the 159 proteins were modeled as linear functions of the proteins for each cell type (linear mixed models, fixed effects: intercept and slope, random effects: intercept and slope). The p values for the slope were then adjusted for both the number of proteins (159) and the number of cell types (14) and thus a total of 2226 tests. RNA and protein were considered as strongly correlated or anticorrelated if the adjusted p < 0.05. Benjamini & Hochberg procedure (Benjamini and Hochberg, 1995) was used for multi-testing correction.

