## Supplementary figures and images for "Multimodal Analysis for Human ex vivo Studies Shows Extensive Molecular Changes from Delays in Blood Processing"

### Savage Figure S1-PBMC metrics, flow cytometry gating

#### Supplemental Figure 1

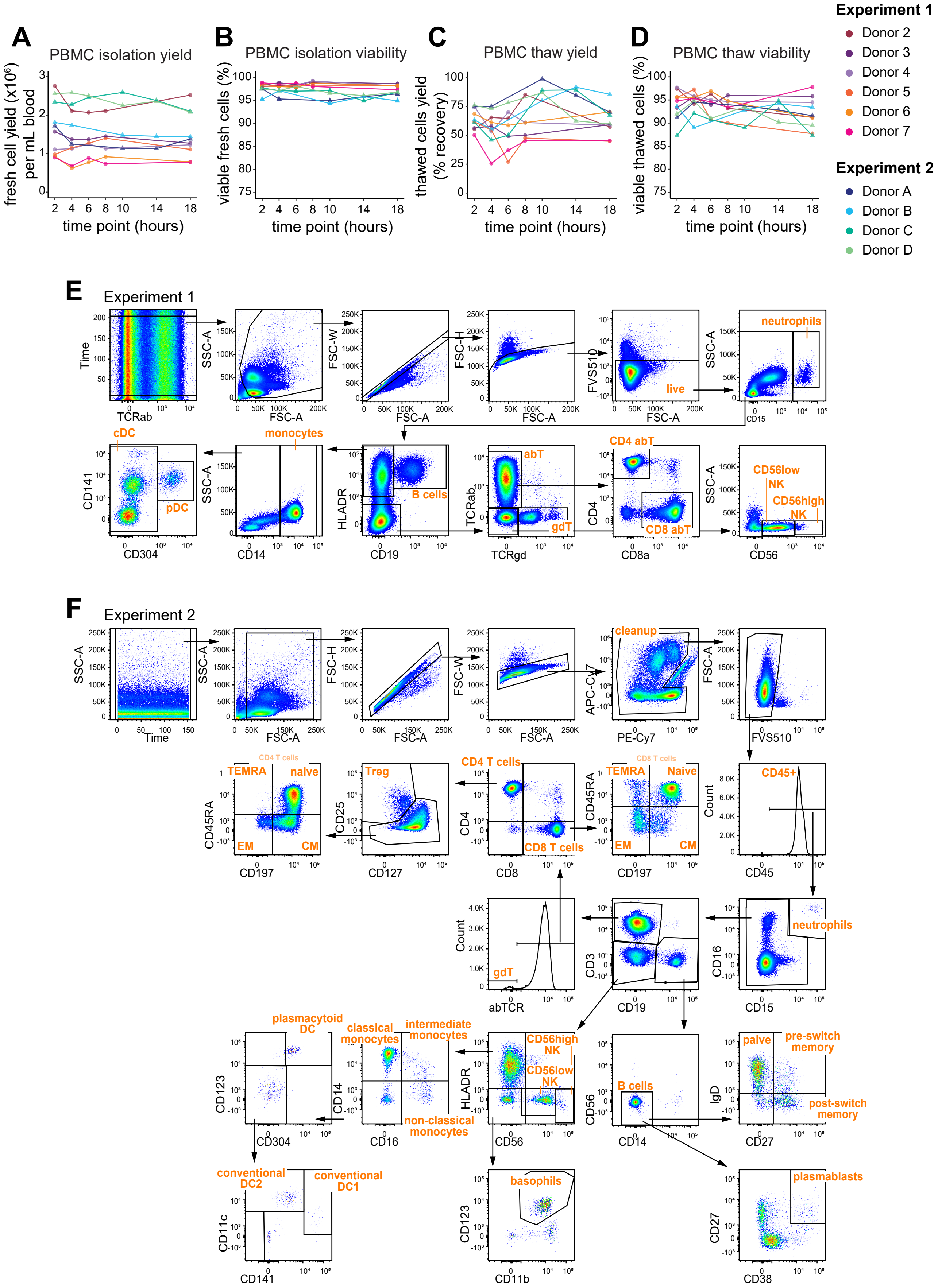

### Savage Figure S2-Flow cytometry frequencies

#### Supplemental Figure 2

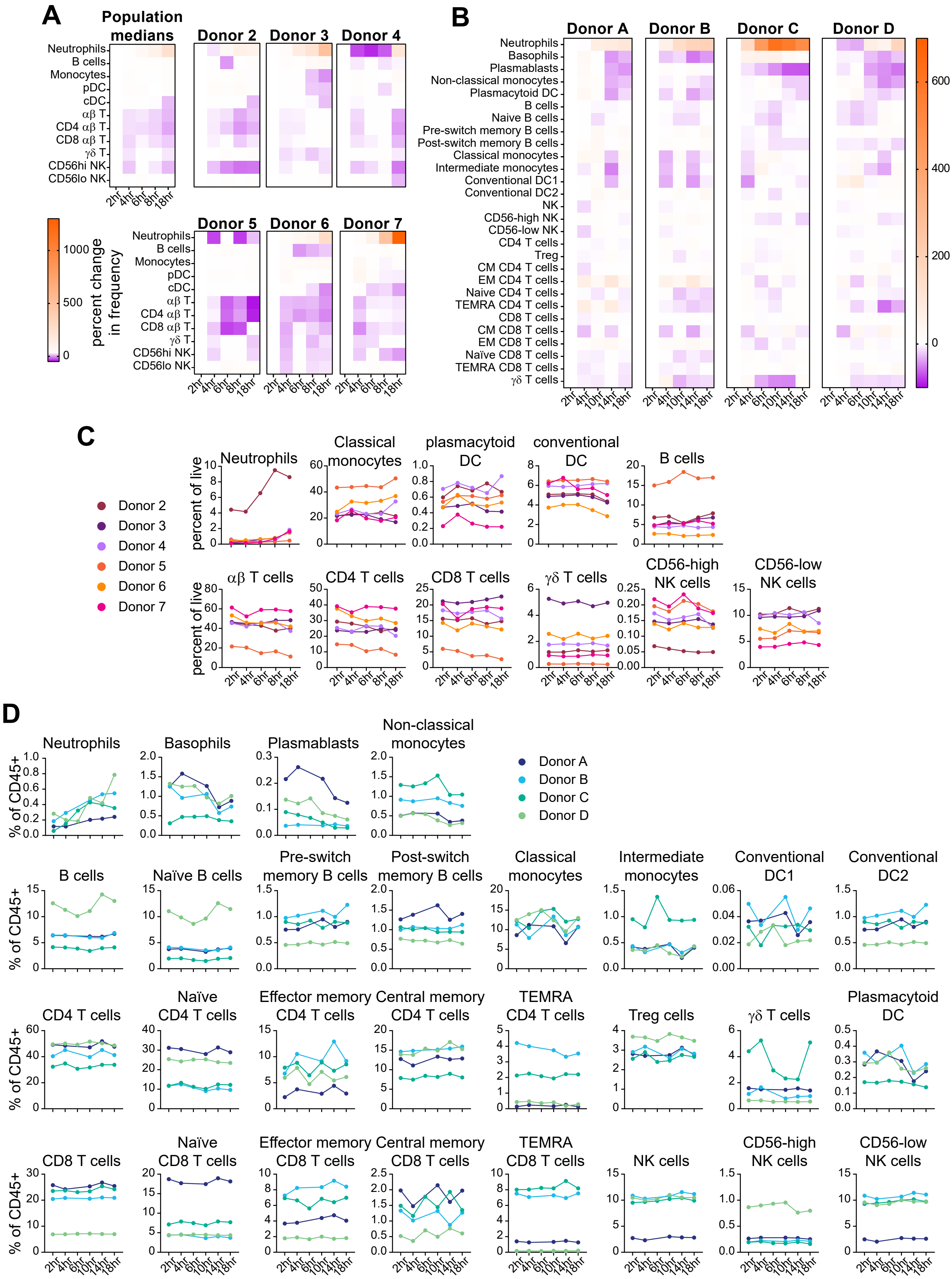

### Savage Figure S3-Nanostring bulk transcriptomics PCA

#### Supplemental Figure 3

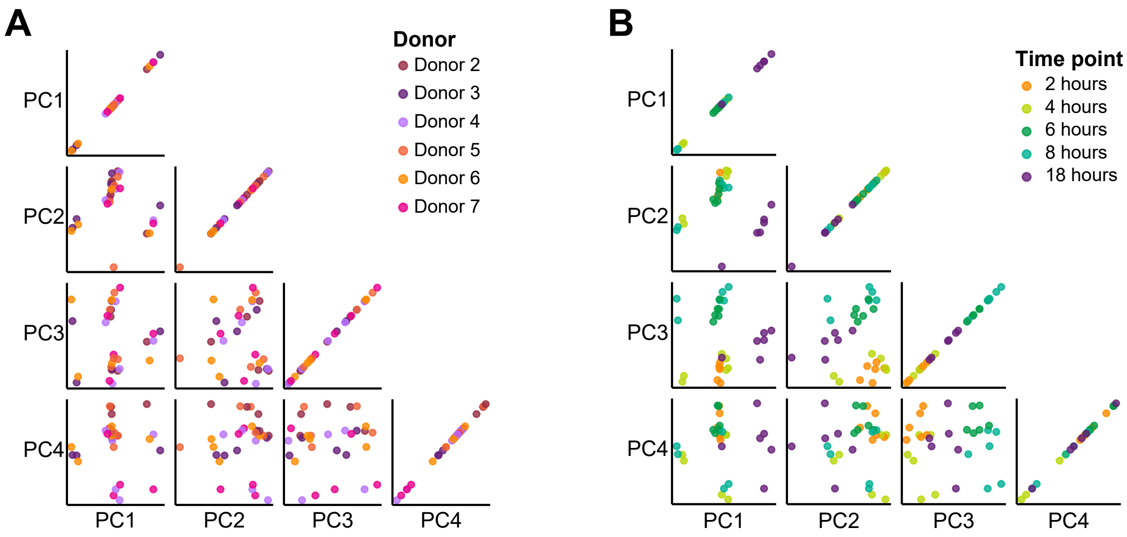

### Savage Figure S4-Single-cell RNAseq, technical metrics

#### Supplemental Figure 4

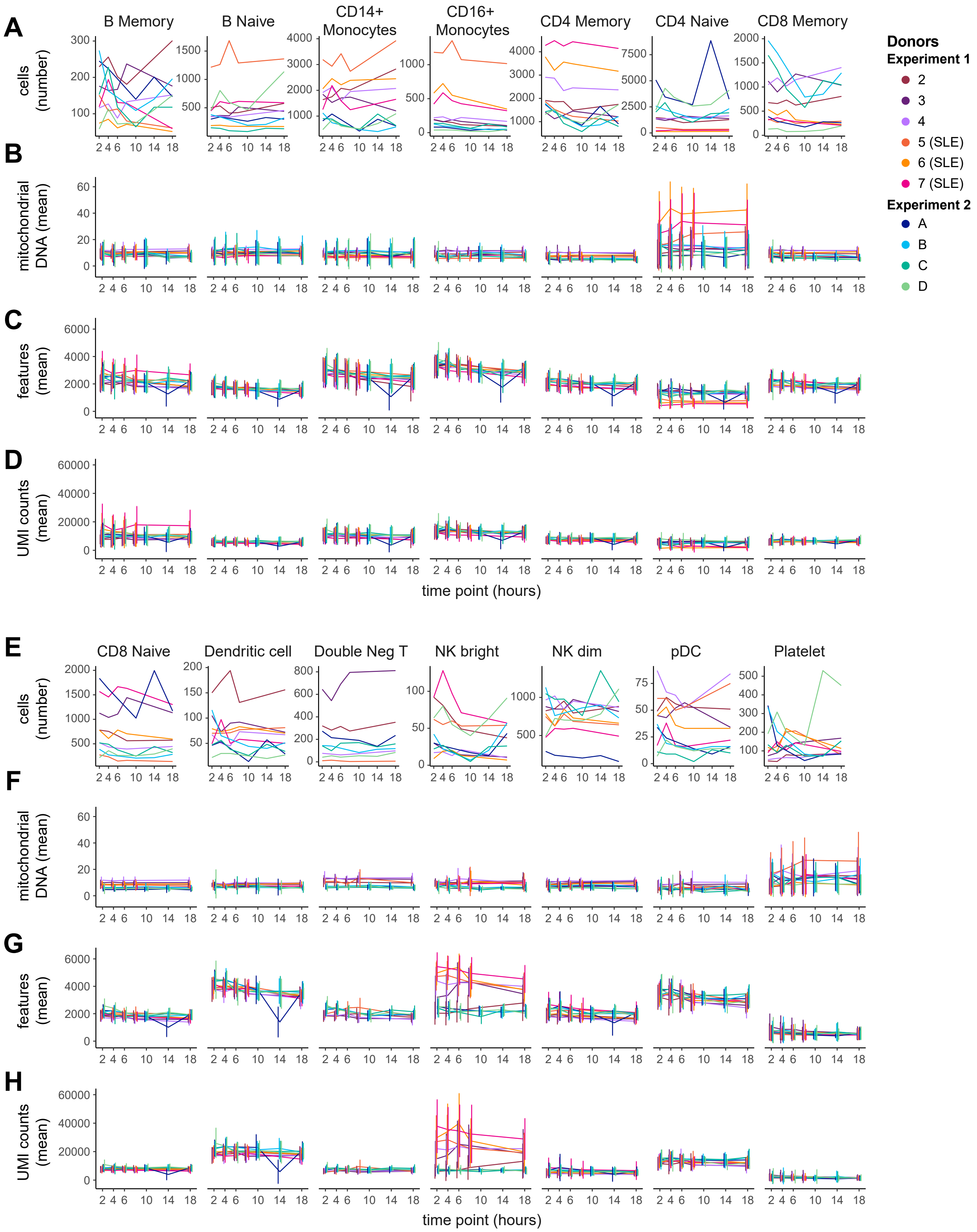

### Savage Figure S5-Single-cell RNAseq, tSNE by time

#### Supplemental Figure 5

**A**

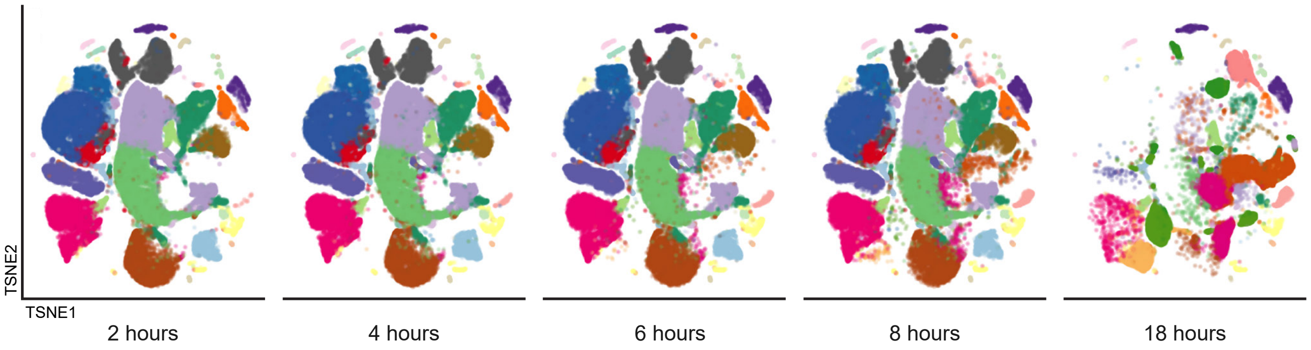


**B**

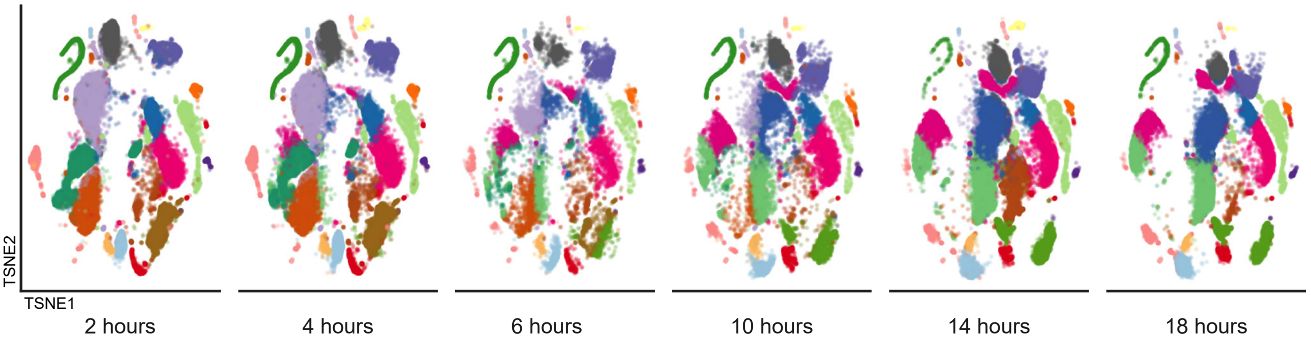

### Savage Figure S7-Protein-transcript correlations

#### Supplemental Figure 7

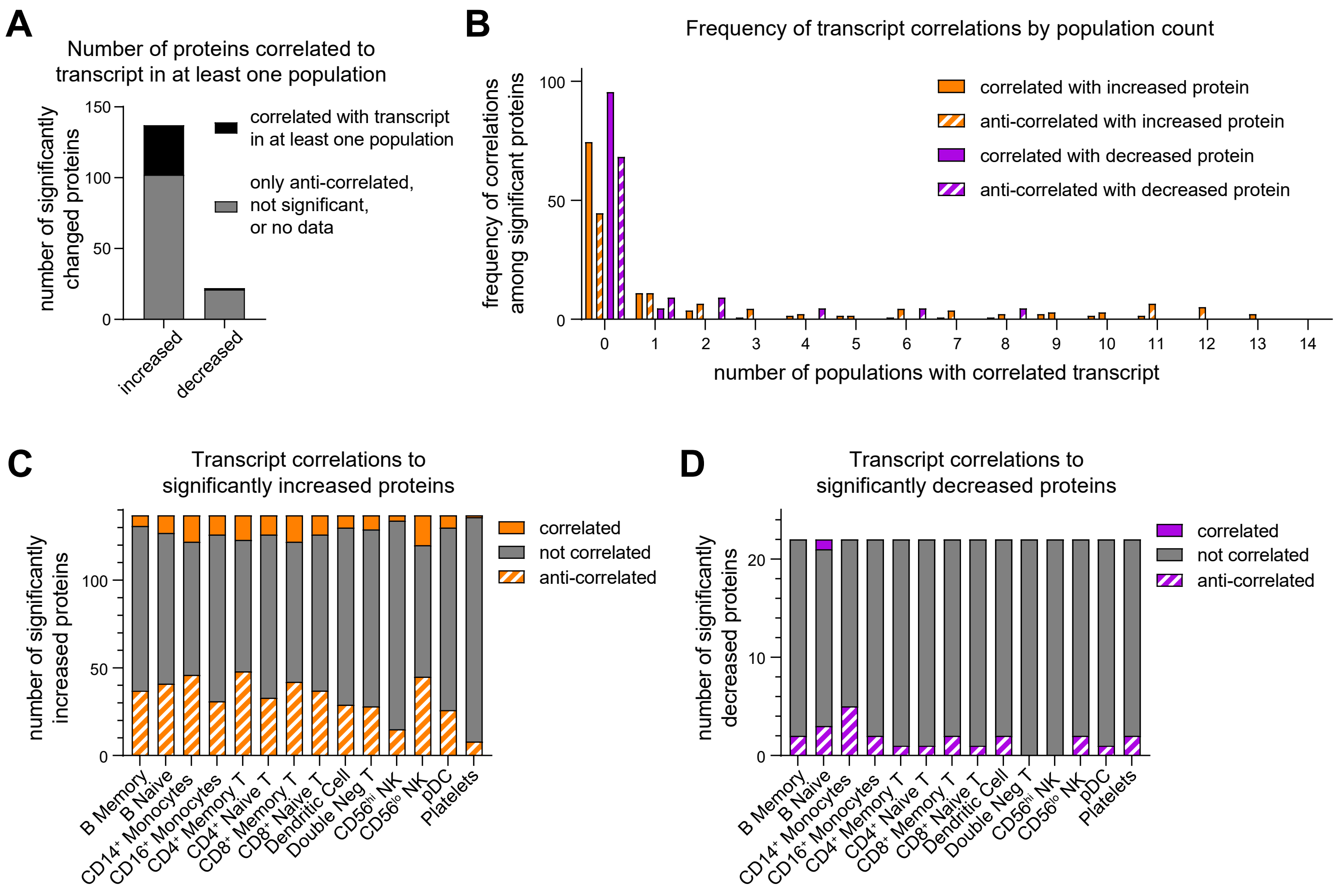

### Savage Figure S8-technical variance

#### Supplemental Figure 8

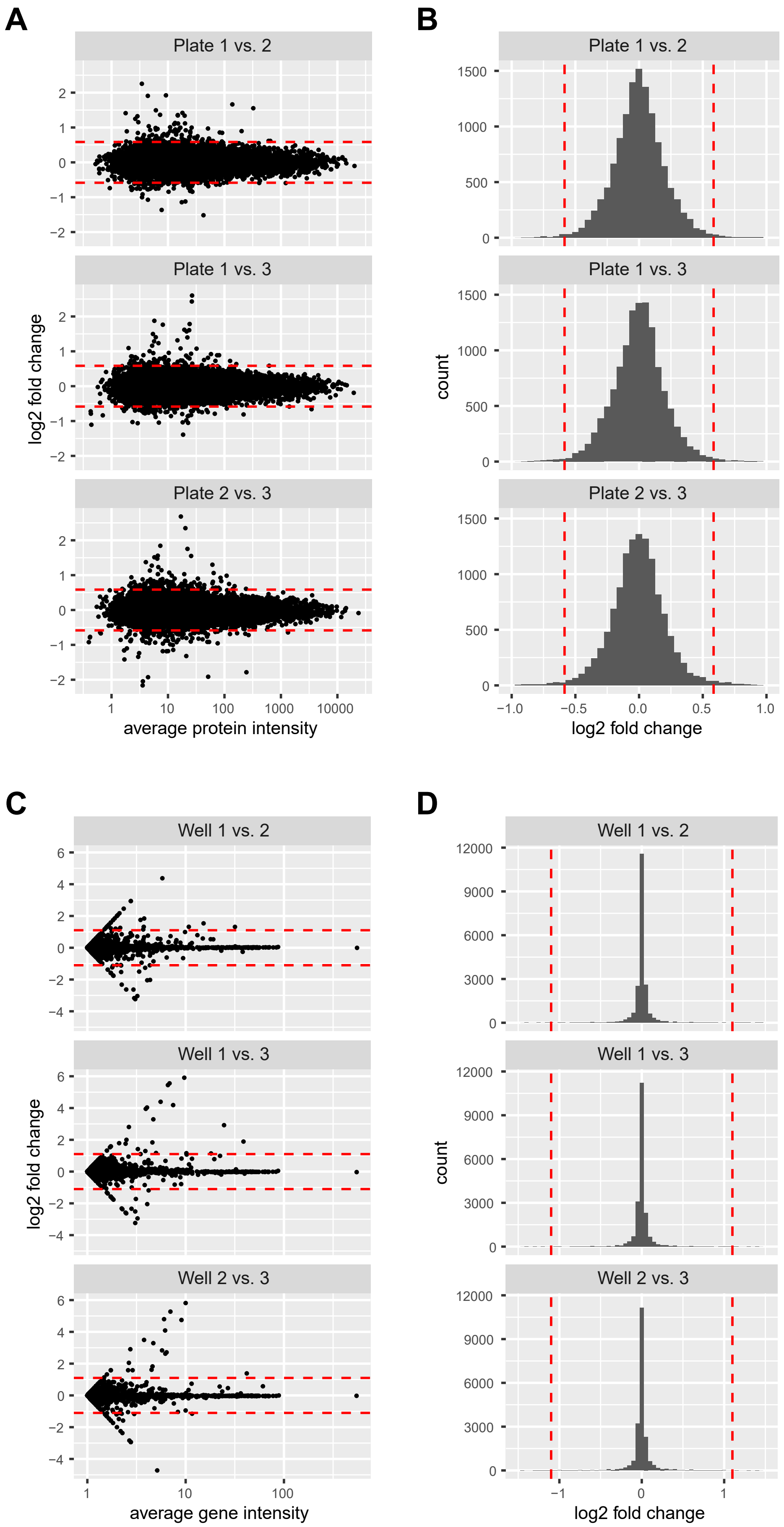
