## Supplementary material for "Multimodal Analysis for Human ex vivo Studies Shows Extensive Molecular Changes from Delays in Blood Processing": Savage Figure S6-Single-cell RNAseq, DEG

### Supplemental Figure 6

**A** Differentially expressed genes - Experiment 1 Heatlhy Donors

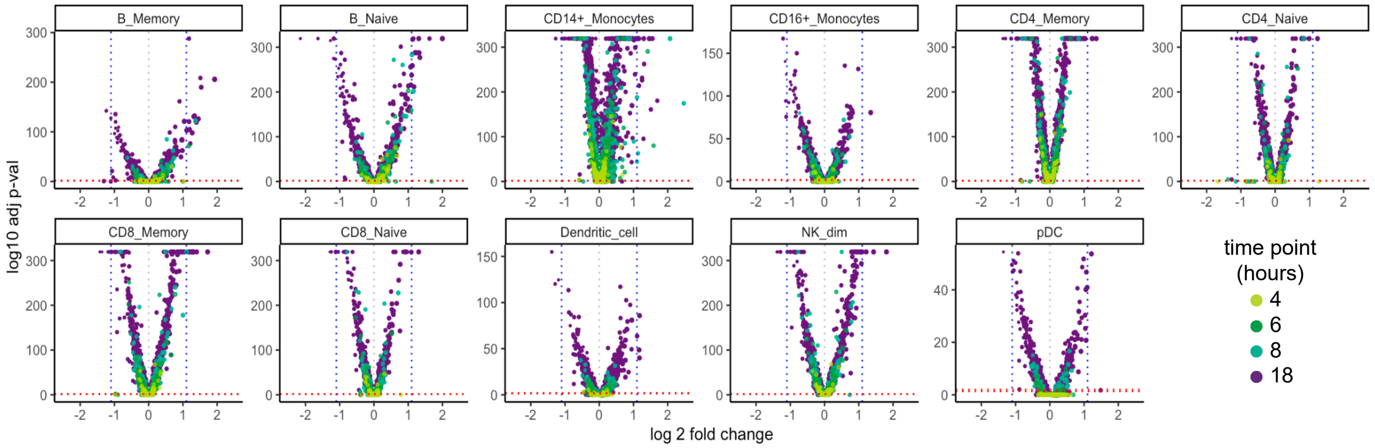


**B** Differentially expressed genes - Experiment 1 SLE Donors

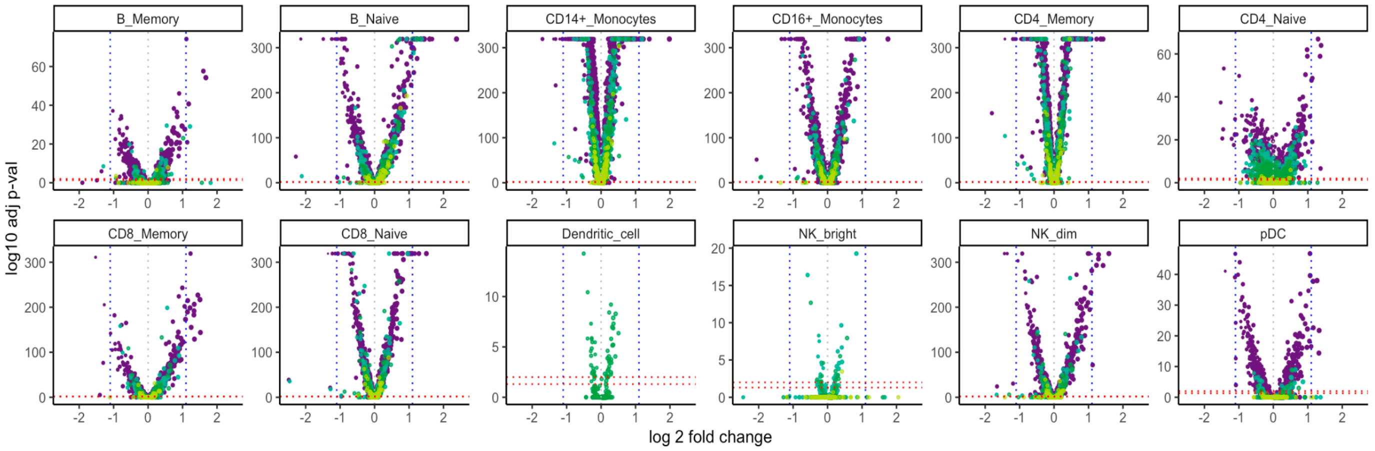


**C** Differentially expressed genes - Experiment 2 Healthy Donors

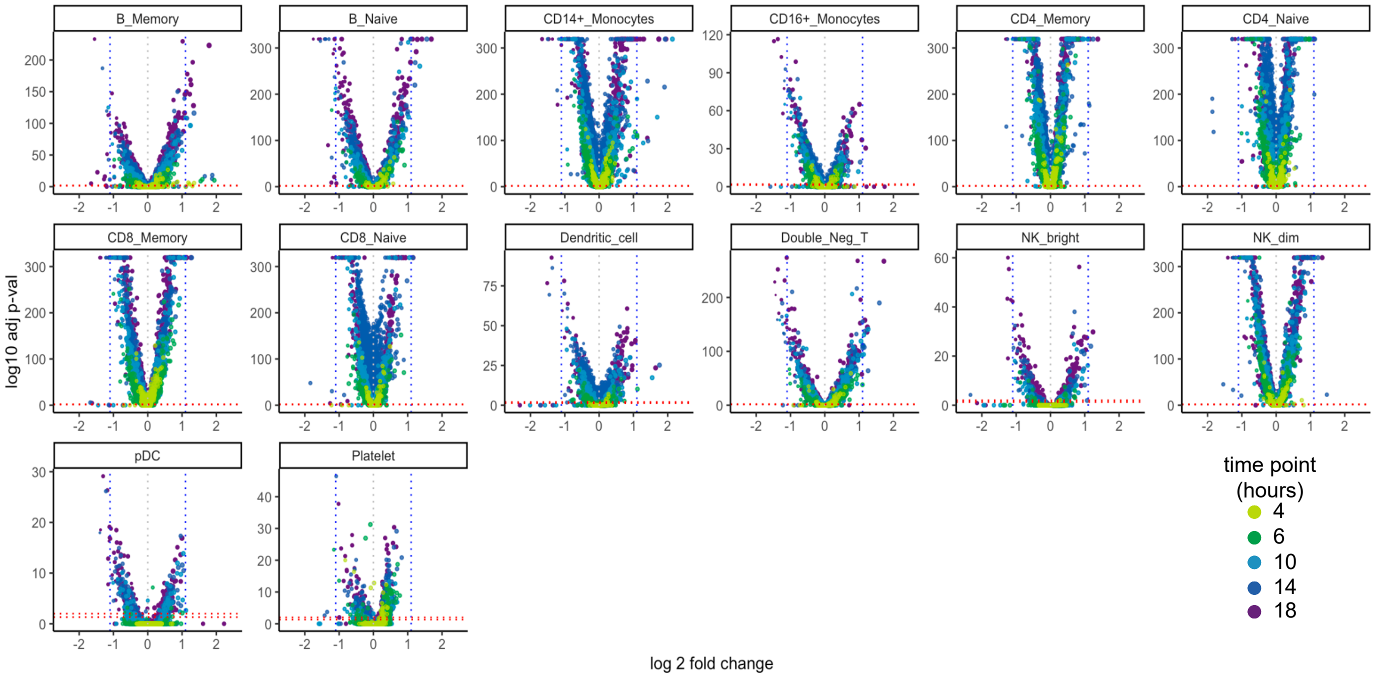
