## Supplementary material for "Multimodal Analysis for Human ex vivo Studies Shows Extensive Molecular Changes from Delays in Blood Processing": Savage Table S7-Donor information

**Experiment 1**

| | | | | | | | |
|---|---|---|---|---|---|---|---|
| | **Sample ID** | 2 | 3 | 4 | 5 | 6 | 7 |
| | **Donor Type** | Healthy | Healthy | Healthy | SLE | SLE | SLE |
| **Demographics** | **Age** | 25 | 25 | 27 | 30 | 25 | 33 |
| | **Sex** | male | male | male | female | female | female |
| | **Race (self-reported)** | White, Caucasian | White, Caucasian | White, Caucasian | White, Caucasian | White, Caucasian, Native Hawaiian, Other Pacific Islander | White, Caucasian |
| | **Hispanic/Latino Ethnicity (self-reported)** | no | no | no | no | no | no |
| **CBC RESULTS** | **Absolute Basophils (cells/uL)** | 47 | 50 | 50 | 22 | 31 | 11 |
| | **Absolute Eosinophils (cells/uL)** | 265 | 99 | 140 | 81 | 220 | 69 |
| | **Absolute Lymphocytes (cells/uL)** | 3050 | 1981 | 1751 | 988 | 1488 | 1341 |
| | **Absolute Monocytes (cells/uL)** | 577 | 341 | 351 | 510 | 604 | 286 |
| | **Absolute Neutrophils (cells/uL)** | 3861 | 4629 | 2210 | 1099 | 3758 | 3593 |
| | **Hematocrit (%)** | 42.3 | 43.1 | 41.1 | 29.5 | 41 | 37 |
| | **Hemoglobin (g/dL)** | 14.6 | 14.8 | 14.6 | 9.7 | 14.1 | 12.2 |
| | **Mch (pg)** | 31.7 | 29.1 | 31.7 | 28.2 | 31.8 | 29.3 |
| | **Mchc (g/dL)** | 34.5 | 34.3 | 35.5 | 32.9 | 34.4 | 33 |
| | **Mcv (fL)** | 92 | 84.7 | 89.3 | 85.8 | 92.6 | 88.9 |
| | **Percent Basophils** | 0.6 | 0.7 | 1.1 | 0.8 | 0.5 | 0.2 |
| | **Percent Eosinophils** | 3.4 | 1.4 | 3.1 | 3 | 3.6 | 1.3 |
| | **Percent Lymphocytes** | 39.1 | 27.9 | 38.9 | 36.6 | 24.4 | 25.3 |
| | **Percent Monocytes** | 7.4 | 4.8 | 7.8 | 18.9 | 9.9 | 5.4 |
| | **Platelet Count (thousand/uL)** | 250 | 316 | 223 | 269 | 279 | 245 |
| | **Rdw Rbc Distribution Width (%)** | 12.2 | 12.7 | 12.3 | 13 | 11.5 | 11.7 |
| | **Red Blood Cell Count (millions/uL)** | 4.6 | 5.09 | 4.6 | 3.44 | 4.43 | 4.16 |
| | **White Blood Cell Count (thousand/uL)** | 7.8 | 7.1 | 4.5 | 2.7 | 6.1 | 5.3 |
| | **Smoking History** | [no data] | [no data] | [no data] | Never Smoker | Former Smoker | Never Smoker |
| | **Duration of SLE (years)** | [not applicable] | [not applicable] | [not applicable] | 8.3 | 5.8 | 4 |
| | **Notes at time of draw** | [none] | [none] | [none] | Stable lupus | Active symptoms - pleurisy, joint stiffness (hip, lower extremities), rash, fatigue | Not flaring - doing well |
| **Treatment #1** | **Medication #1 Name** | [not applicable] | [not applicable] | [not applicable] | Hydroxychloroquine | Hydroxychloroquine | Hydroxychloroquine |
| | **Dosage** | [not applicable] | [not applicable] | [not applicable] | 200 mg | 200 mg | 200 mg |
| | **Frequency** | [not applicable] | [not applicable] | [not applicable] | 2x a day | 2x a day | 2x a day |
| | **Duration of treatment** | [not applicable] | [not applicable] | [not applicable] | 8 years | 5.65 years | 3.87 years |
| **Treatment #2** | **Medication #2 Name** | [not applicable] | [not applicable] | [not applicable] | [none] | Methotrexate | [none] |
| | **Dosage** | [not applicable] | [not applicable] | [not applicable] | [none] | 7.5 mg | [none] |
| | **Frequency** | [not applicable] | [not applicable] | [not applicable] | [none] | 1x a week | [none] |
| | **Duration of treatment** | [not applicable] | [not applicable] | [not applicable] | [none] | 7 months | [none] |

### Table S7

**Experiment 2**

| Sample ID | A | B | C | D |
|---|---|---|---|---|
| **Donor Type** | Healthy | Healthy | Healthy | Healthy |
| **Age** | 33 | 51 | 26 | 51 |
| **Sex** | female | male | male | female |
| **Race (self-reported)** | [no answer] | [no answer] | [no answer] | White |
| **Hispanic/Latino Ethnicity (self-reported)** | Caucasian | Caucasian | White | Non-hispanic |
| **Smoking status** | no | no | no | no |
| **LYM (%)** | 28.2 | 33.1 | 36.3 | 33.4 |
| **LYM (10^3/mm3)** | 1.6 | 1.9 | 2.3 | 2.6 |
| **MON (%)** | 6 | 4.8 | 5.3 | 4.6 |
| **MON (10^3/mm3)** | 0.3 | 0.2 | 0.3 | 0.3 |
| **GRA (%)** | 65.8 | 62.1 | 58.4 | 62 |
| **GRA (10^3/mm3)** | 3.9 | 3.8 | 3.9 | 5.1 |
| **WBC (10^3/mm3)** | 5.8 | 5.9 | 6.5 | 8 |
| **RBC (10^6/mm3)** | 3.9 | 4.9 | 4.5 | 4.8 |
| **HGB (g/dL)** | 12.6 | 14.8 | 14.7 | 13.2 |
| **HCT (%)** | 37.8 | 44.4 | 43.1 | 41.4 |
| **PLT (10^3/mm3)** | 287 | 232 | 237 | 351 |
| **MCV (um3)** | 96 | 91 | 95 | 86 |
| **MCH (pg)** | 31.9 | 30.3 | 32.5 | 27.3 |
| **MCHC (g/dL)** | 33.2 | 33.4 | 34.2 | 32 |
| **RDW (%)** | 13.5 | 13.5 | 13 | 16 |
| **MPV (um3)** | 7 | 7.8 | 7.7 | 6.7 |
